# Passage of the HIV capsid cracks the nuclear pore

**DOI:** 10.1101/2024.04.23.590733

**Authors:** Jan Philipp Kreysing, Maziar Heidari, Vojtech Zila, Sergio Cruz-Leon, Agnieszka Obarska-Kosinska, Vibor Laketa, Sonja Welsch, Jürgen Köfinger, Beata Turoňová, Gerhard Hummer, Hans-Georg Kräusslich, Martin Beck

## Abstract

Upon infection, human immunodeficiency virus (HIV-1) releases its cone-shaped capsid into the cytoplasm of infected T-cells and macrophages. As its largest known cargo, the capsid enters the nuclear pore complex (NPC), driven by interactions with numerous FG-repeat nucleoporins (FG-Nups). Whether NPCs structurally adapt to capsid passage and whether capsids are modified during passage remains unknown, however. Here, we combined super-resolution and correlative microscopy with cryo electron tomography and molecular simulations to study nuclear entry of HIV-1 capsids in primary human macrophages. We found that cytosolically bound cyclophilin A is stripped off capsids entering the NPC, and the capsid hexagonal lattice remains largely intact inside and beyond the central channel. Strikingly, the NPC scaffold rings frequently crack during capsid passage, consistent with computer simulations indicating the need for NPC widening. The unique cone shape of the HIV-1 capsid facilitates its entry into NPCs and helps to crack their rings.

## Introduction

Human immunodeficiency virus type 1 (HIV-1) is a lentivirus that can infect non-dividing cells including human macrophages ^1^. Fusion of the viral and cellular membrane leads to cytoplasmic entry of the characteristic cone-shaped HIV-1 capsid that encases the viral genome and replication machinery. Subsequently, the viral genomic RNA undergoes reverse transcription into double-stranded cDNA, which eventually becomes integrated into the host cell genome, where it is maintained for the life of the infected cell. Reverse transcription initiates in the cytoplasm and is completed after nuclear entry of the subviral replication complex prior to integration into the host cell genome ^2-4^. HIV-1 subviral complexes comprise the viral genome associated with nucleocapsid proteins and the replication proteins reverse transcriptase (RT) and integrase (IN). While earlier studies indicated rapid disassembly of the capsid in the cytoplasm releasing the free replication complex, it is now clear that the capsid can be stably maintained up to nuclear entry ^5, 6^. During cytoplasmic transit, the HIV-1 capsid has been shown to engage multiple host cell restriction and dependency factors, to serve as a reaction container for reverse transcription and to shield the nascent viral cDNA from cytoplasmic antiviral DNA sensors ^5, 6^. Furthermore, the capsid lattice engages microtubular motors, thus facilitating its transport towards the nuclear envelope ^7-9^. Capsid also interacts with the cytoplasmic protein cyclophilin A (CypA) ^10, 11^ and with the Cyp domain of Nup358 at the cytoplasmic side of nuclear pores, potentially docking the capsid to the nuclear pore complex (NPC) ^12^. Accordingly, the HIV-1 capsid is the main orchestrator of the early post-entry phase of viral replication.

A potential role of the HIV-1 capsid for nuclear entry and in the nucleoplasm of non-dividing cells has long been under debate ^5, 6^.The capsid consists of 200-250 hexamers of the viral CA protein with 12 CA pentamers inserted in regions of high curvature ^13-15^. The hexamers have been shown to specifically interact with phenylalanine-glycine (FG) repeats that are present within intrinsically disordered regions of several nucleoporins (FG-Nups) ^16, 17^.The FG-binding hydrophobic cleft within the CA hexamer also interacts with an FG-containing motif of the nuclear protein cleavage and polyadenylation specificity factor 6 (CPSF6) ^17^. FG-motif containing host factors can be displaced by small molecules (e.g., Lenacapavir ^18^), competitively binding the CA pocket ^19^. Lenacapavir has recently been approved for treatment of HIV-1 infected patients. The capsid-targeting compounds can block nuclear entry of HIV-1 replication complexes, supporting a role of the capsid during NPC passage ^16, 19, 20^. The size discrepancy between the HIV-1 capsid (ca. 60 nm at the broad end ^15^) and the inner diameter of NPCs obtained from isolated nuclear envelopes (ca. 45 nm ^21^) had led to the conclusion that genome uncoating must occur prior to entry into the NPC channel, possibly leaving some CA remnants on the subviral complex. This hypothesis was supported by the observation that a CypA-DsRed fluorescent fusion protein, which efficiently binds cytoplasmic capsids, was rapidly lost when the fluorescent structure reached the nuclear pore ^22^. Recent ultrastructural studies have shown, however, that (largely) intact capsids can enter the nucleus in a T-cell line through apparently normal nuclear pores ^23^, and cone-shaped capsids encasing electron-dense nucleoprotein complexes have been detected inside the nucleus of HIV-1 infected reporter cell lines and primary human macrophages ^23,4,24^. This was explained by the recent observation, that the central channel of NPCs can adapt its diameter in response to forces laterally imposed by the nuclear membranes ^25^. With an inner diameter of approximately 45-65 nm ^23, 26-28^ the NPC central channel of human tissue culture cells appeared to be sufficiently wide for passage of the complete HIV-1 capsid.

NPCs consist of three sandwiched rings. The nuclear (NR), inner (IR) and cytoplasmic ring (CR) are 8-fold rotationally symmetric structures that form a central channel thus bridging across the inner and outer membranes of the nuclear envelope. The conformational changes that occur during NPC diameter changes differ markedly between the individual rings. While the CR and NR form an elaborate and thoroughly connected scaffold that is bent during dilation movements, the IR consists of eight spokes that are flexibly connected. These spokes move outwards and away from each other during NPC dilation, thereby stretching their linkers and opening up additional space within the central channel ^26, 29^. If and how the dilation of individual NPC is coordinated with the transport of large cargos remains unknown, however.

Nuclear transport cargo passes through the central channel, which is filled with tens of megadaltons of intrinsically disordered FG-Nups. These FG-Nups selectively bind cargo to facilitate nuclear import, but at the same time exclude the vast majority of proteins from entering the nucleus ^30^. FG-Nups thus form the permeability barrier for the nuclear envelope, protect the host cell genome and constitute a major barrier that would have to be overcome by the HIV-1 capsid for nuclear entry ^31^. However, two recent studies showed that in vitro assembled HIV-1 capsids rapidly partitioned into phase separated condensates of the intrinsically disordered FG repeat region of Nup98, and partitioning was dependent on the FG binding pocket in CA ^32, 33^. These results confirmed that the viral capsid itself constitutes a multivalent nuclear import cargo with surface properties similar to nuclear transport receptors. Upon reaching the nucleus, the capsid must eventually break open to release the reverse transcribed genome for integration to occur. Although mounting evidence indicates that transcriptional latency, as the main obstacle for HIV-1 cure, is mediated in part by the position of genome integration ^34, 35^, neither the mechanism nor the regulation of time or place of uncoating are currently understood.

The two central questions regarding nuclear import of the HIV-1 capsid are: How do capsids pass through a channel of comparable width that is densely filled with FG-Nups? And further, at what location do capsids open to release the genome? Three scenarios may be considered, which are not mutually exclusive. (i) The capsid lattice may be elastic, a hypothesis that is supported by a recent preprint reporting the capsid lattice to undergo reversible deformation without disintegration ^36^. (ii) The capsid lattice may break during NPC passage, thus altering its geometry. While morphologically normal appearing capsids have been detected in the nucleoplasm of non-dividing cells, lattice analysis so far has been performed only on cell-free virions and lattice completeness of intracellular capsids has not been determined. (iii) Passage of the HIV-1 capsid through the NPC channel may induce local NPC dilation, possibly even to a point where the NPC scaffold is altered, thereby giving room for intact capsids to pass.

Here, we provide direct evidence for loss of CypA from HIV-1 capsids at the nuclear pore. We show that NPC passage appears rate-limiting in primary human macrophages and we identify apparently intact, nucleic acid-containing capsids in the nucleoplasm of these cells. Most importantly, we provide experimental evidence that passage of intact HIV-1 capsids cracks the NPC architecture. Computational simulation supports a scenario in which cracking of the NPC scaffold facilitates capsid passage by relieving a steric barrier.

## Results

### Subviral HIV-1 complexes accumulate at nuclear pores in human primary macrophages

We studied nuclear entry of subviral HIV-1 complexes in primary human monocyte-derived macrophages (MDM). Macrophages are natural target cells of HIV-1 and are post-mitotic; accordingly, the viral replication complex has to enter the nucleus through the intact nuclear envelope. To analyze this process, we made use of our previously described non-infectious HIV-1 variant (NNHIV) that carries a complete HIV-1 genome with deleterious mutations in IN and a deletion in the *tat* gene. We have previously shown that NNHIV can enter permissive cells and undergo reverse transcription and nuclear entry similar to wild-type HIV-1, but is defective in integration and gene expression ^23^.

Freshly prepared MDM were incubated with NNHIV carrying a macrophage-tropic HIV-1 envelope (Env) glycoprotein for different periods of time and fixed cells were immunostained for HIV-1 CA and for lamin A/C to define the nuclear boundary. Using confocal microscopy, we observed significant variability in efficiency of subviral complexes to reach the nucleus between individual cells from one donor as well as between different donors. Some cells exhibited CA signals distributed through the whole cell including nuclear regions, while others did not show any signal in the proximity of or inside the nucleus. Focusing on cells displaying intranuclear HIV-1 signals, we observed a strong enrichment of subviral complexes overlaying the lamin signal and thus likely localized at the nuclear envelope. This phenotype was clearly detected at 24 h post infection (p.i.) and was stronger at 48 h p.i. (Figure 1A-C), while CA signals were cytoplasmically distributed at earlier time points. Starting at 24 h p.i., we observed clear HIV-1 signals in the nucleus of NNHIV-treated MDM and their number increased at 48 h p.i. (Figure 1A-C).

**Figure 1.**
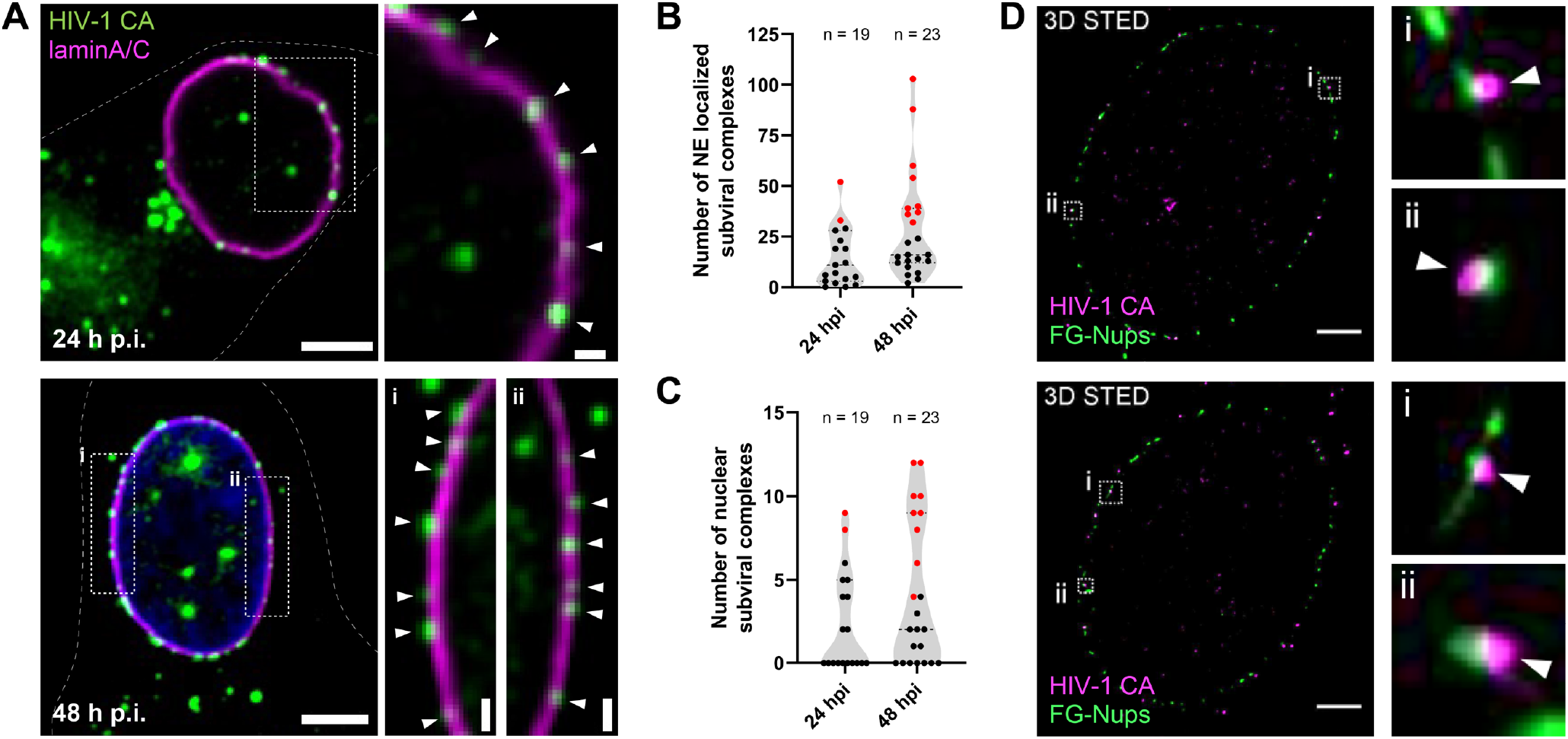
HIV-1 capsids accumulate at nuclear pores in primary human macrophages. (A–C) MDM were infected with macrophage-tropic NNHIV for 24 h (A, upper) or 48 h (A, lower). Cells were then fixed, methanol extracted and immunostained for HIV-1 CA (green) and laminA/C (magenta). Shown are confocal z slices through the nuclei of infected MDM. The enlarged regions (A, right) display HIV-1 CA signals (white arrowheads) accumulated at the nuclear envelope (NE) as defined by laminA/C staining. (B, C) Quantification of CA signals representing subviral complexes co-localizing with laminA/C (B) and number of CA signals detected inside the nucleus (C) at indicated times p.i. Indicated in red are values from cells exhibiting >30 CA signals colocalizing with laminA/C. Results represent data obtained by analysis of (n) cells from one representative donor. (D) 3D STED imaging of nuclei of HIV-1 infected MDM. Cells were infected for 48 h, fixed, methanol extracted and immunostained for HIV-1 CA (magenta) and FG-Nups (green). Slices through the 3D reconstruction of an entire nucleus are shown with enlargements (D, right) displaying CA signals (white arrowheads) directly associated with NPCs as defined by FG-Nup staining. See also Video S1. Scale bars: 2 μm.

To analyze whether nuclear envelope-associated capsids were enriched at NPCs, we performed stimulated emission depletion (STED) nanoscopy of primary MDM from different donors. Cells were incubated with NNHIV for 48 h and stained for HIV-1 CA and FG-Nups. Subsequently, we performed two-color 3D STED nanoscopy and acquired super-resolved images in sequential optical sections to cover the entire nuclear volume of individual cells. Analysis of computational slices of fully reconstructed MDM nuclei confirmed the direct association of the HIV-1 capsid with nuclear pores (Figure 1D, Video S1). Quantitative analysis showed that 89% (908 of 1,025) of all CA signals detected at the nuclear envelope were closely associated with nuclear pores. These results clearly showed that HIV-1 capsids specifically associate with nuclear pores in infected MDM, and their strong accumulation suggests that passage through the nuclear pore is a rate-limiting event in early HIV-1 replication.

### HIV-1 capsids are detected upon entry into, passage through and exit from the NPC

In order to determine the relative positioning of HIV-1 capsids at the nuclear pore, we combined two-color STED nanoscopy for CA and FG-Nups with nuclear Hoechst staining in confocal mode to define the nuclear side. Maximum intensity of the FG repeat immunofluorescence signal is expected to localize to the central channel at the inner ring of the nuclear pore ^37^. Analyzing cells at 48 h p.i., we observed CA signals in three positions: at the cytoplasmic side, directly overlapping the FG signal and at the nucleoplasmic side of the nuclear pore (Figure 2A). For quantitative analysis, we segmented individual capsid-associated pores and determined the normalized signal intensities of the FG-Nup and CA signals for a line profile (Figure 2B). This analysis confirmed that HIV-1 capsids could be observed upon their entry into, passage through and exit from the central channel of the NPC in infected MDM. The distribution for these three different positions relative to the center of the nuclear pore was determined for a total of 180 capsid-associated nuclear pores in MDM from three different donors (Figure 2C). A similar distribution was observed for all three donors with 51 % of capsids observed on the cytoplasmic side of the nuclear pore, 33 % overlapping the FG repeat in the central channel and 15 % on the nucleoplasmic side (Figure 2C). These results indicated that entry into and passage through the nuclear pore may be delaying or limiting nuclear entry of HIV-1 capsids in MDM. The substantial number of capsids observed at the nucleoplasmic side of NPCs further suggested that release from the nuclear basket may not be instantaneous after passage through the central channel.

**Figure 2.**
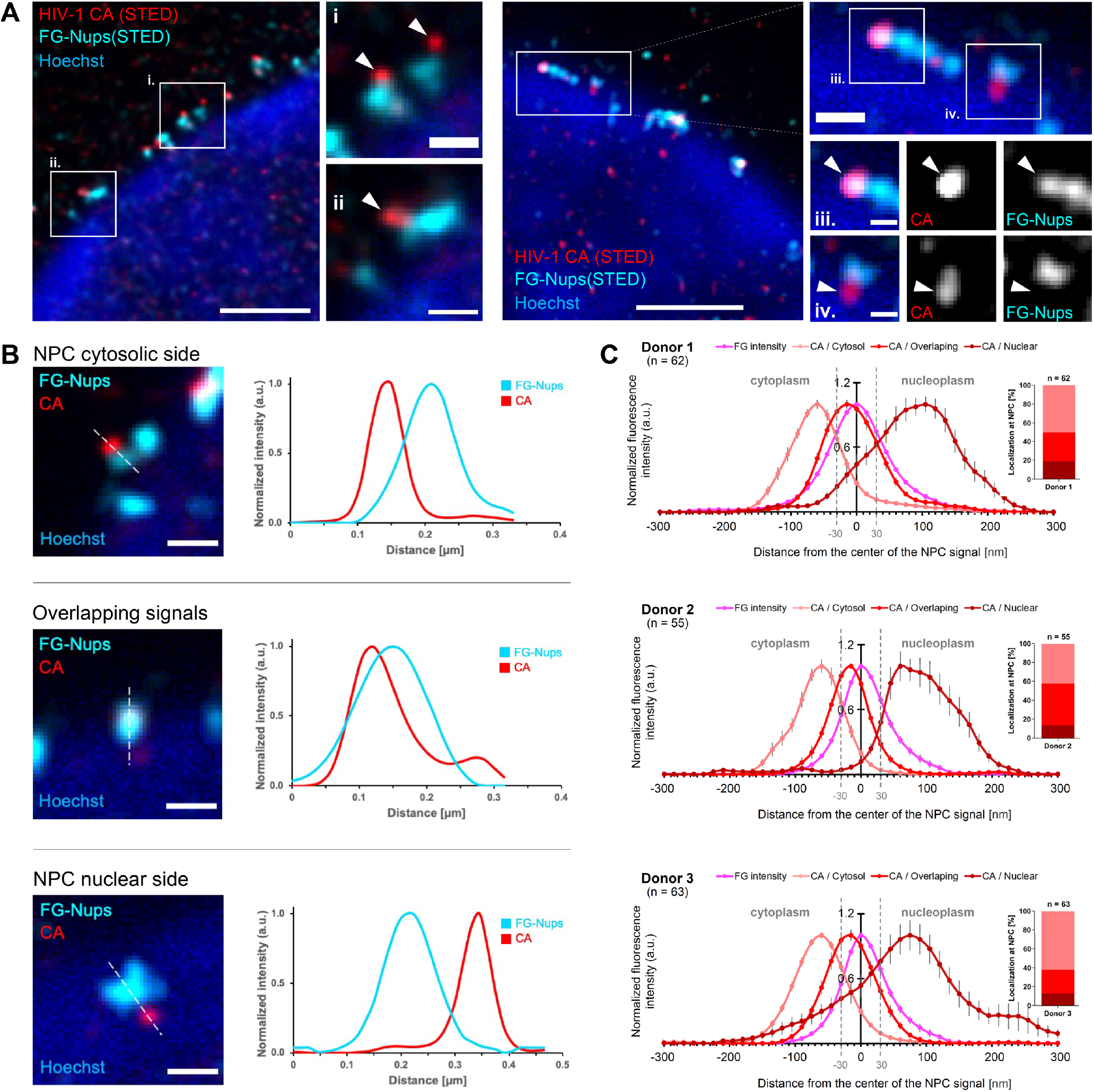
Super resolution analysis of HIV-1 subviral complexes at nuclear pores of infected MDM. MDM were infected with NNHIV-1, fixed at 48 h post-infection and immunostained for HIV-1 CA (red) and FG-Nups (cyan). The nuclear region was visualized by staining with Hoechst and analysed in confocal mode (blue). (A) STED microscopy images displaying nuclear segments of two infected MDM. The enlarged regions show CA signals (white arrowheads) at the cytoplasmic side of the NPC (i. and ii.), overlapping CA and NPC signals (iii.), and CA signals located at the nuclear side of the NPC (iv.). (B) Three typical localizations of HIV signals at the NPC with corresponding CA and FG-Nups signal intensity profiles. Graphs show intensities normalized to the respective maximal value measured in line profiles indicated in images on the left. (C) Averaged line profiles measured on (n) nuclear pores with associated CA signals from three independent experiments, each using cells from a different donor. Error bars represent SD. The localization of CA signals at the NPC was classified as cytoplasmic side of the NPC (pink line), overlapping with NPC (red line) or nuclear side of the NPC (dark red line), applying a window of 30 nm distance of the CA intensity peak from the FG-Nups (magenta line) intensity peak (dashed vertical lines). Scale bars: 500 nm.

### Subviral complexes passing through the central NPC channel are cone-shaped capsids

The fluorescence imaging results clearly revealed the accumulation of CA containing subviral complexes at and inside nuclear pores of infected MDM, but did not provide sufficient resolution to define the morphology and structure of the cargo. We therefore applied 3D correlative light and electron microscopy (CLEM). MDM were infected with NNHIV carrying a fluorescent fusion protein of the viral IN with mScarlet to identify intracellular subviral HIV-1 complexes. MDM were cryo-immobilized by high pressure freezing at 48 h p.i. and further processed for CLEM. Tilt series were acquired for 61 positions of correlated regions targeting IN.mScarlet signals at nuclear envelope regions (defined by Hoechst staining of resin sections) identified in 17 cells from two donors. From this data set, we identified a total of 43 structures completely covered in the resin sections that resembled HIV-1 capsids inside or immediately adjacent to nuclear pores. Their distribution across the NPC was similar to that observed by STED nanoscopy with 19 subviral particles on the cytoplasmic side, 13 deep inside the NPC and 11 on the nucleoplasmic side (Figure S1). Overall, the morphology of the observed subviral structures closely matched that of mature capsids inside purified HIV-1 particles, including the presence of dense material inside capsid structures indicating the presence of condensed ribonucleoprotein or reverse transcription intermediates (Figure 3 and S1). The majority of structures was cone-shaped (41/43; 95%) with rare tubular structures (2/43; 4.7%) (Figure S1). Capsid-like structures at or within the nuclear pore exhibited an average length of 111 ± 11 nm and an average width of 53 ± 6 nm, similar to the dimensions determined for mature HIV-1 capsids by cryo ET ^38^, and almost identical to the dimensions observed in our previous CLEM analysis of an infected T-cell line ^23^. Importantly, we did not observe apparently empty (lacking dense inner material) or obviously broken capsid-like particles inside or directly associated with the nuclear pore in infected MDM (Fig. S1).

**Figure 3.**
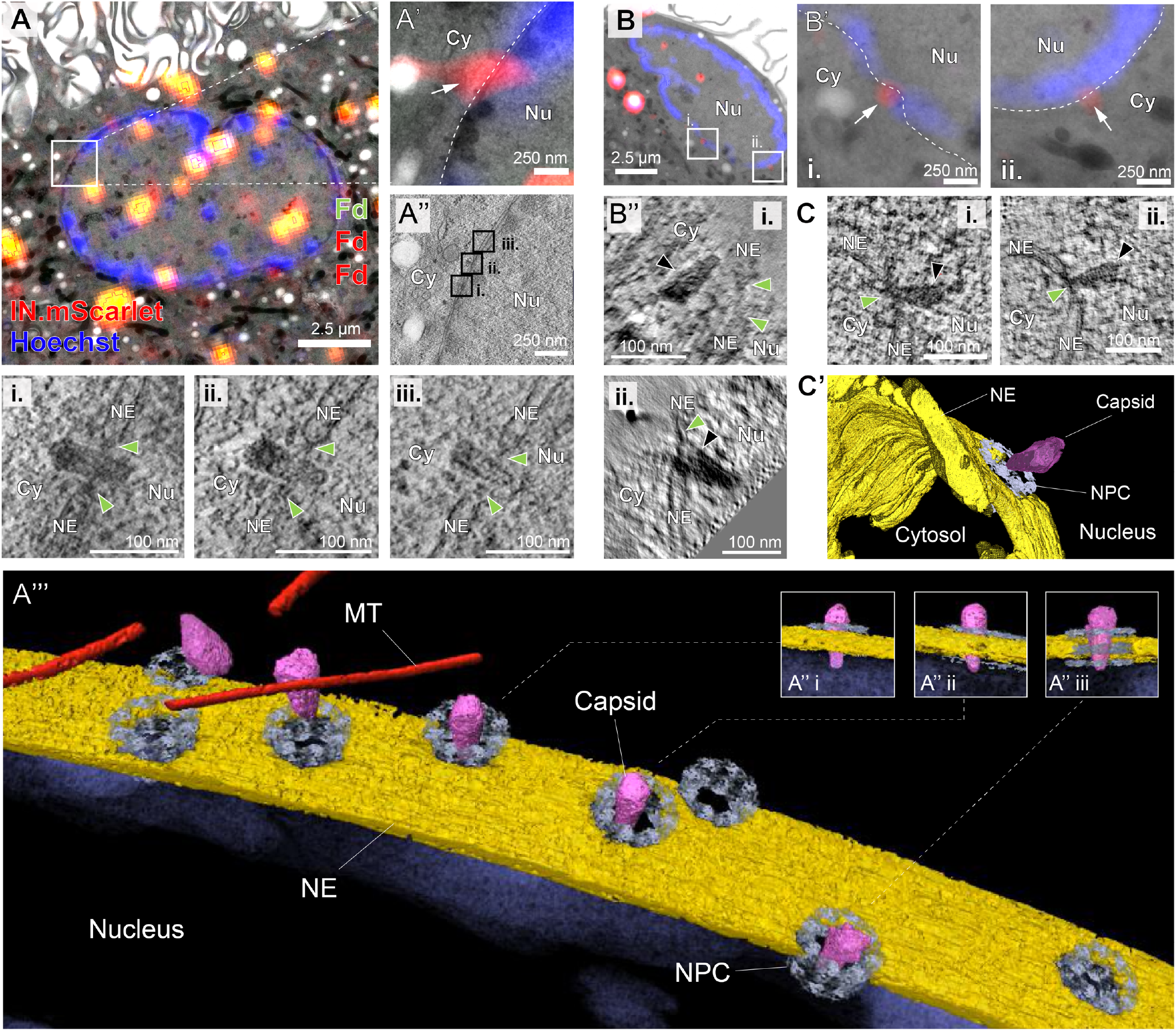
Morphologically intact HIV-1 capsids at and inside NPCs of human MDM. MDM were infected with macrophage-tropic IN.mScarlet carrying NNHIV for 48 h at 37°C, prior to cryo-immobilization by high pressure freezing, embedding and further processing for CLEM-ET. Fluorescently labelled subviral structures in the region of the nuclear envelope were visualized by CLEM-ET. Dashed lines in enlargements outline the nuclear envelope. (A and A′) CLEM overlay (A) with enlargement (A′) showing positions of IN.mScarlet signals (red; white arrow) at the nuclear envelope in an EM section post-stained with Hoechst (blue) and decorated with multi-fluorescent fiducials (Fd) for correlation. (A″) Slice through a tomographic reconstruction at the correlated position shown in (A) and (A′). The features i–iii that are shown enlarged in the bottom panel are framed in black and contain three different capsids that deeply penetrate the central channel of the NPC (green arrowheads). Cy, cytosol; Nu, nucleus; NE, nuclear envelope; NPC, nuclear pore complex. (A′′′) Same as in (A′′) but displayed segmented and isosurface rendered. MT (microtubule) red; capsid, magenta; NE, yellow; NPC, cyan (cryo-EM map of NPC: ^23^). See also Video S2. (B – B′′) Same as (A and A′′) showing capsid docking at the NPC (i) and capsid located in the nuclear basket region (ii) from the same resin section. Both capsids display a conical shape and a dense interior (black arrowheads). (C) Slices through tomographic reconstructions from two different resin sections, showing two cone-shaped capsids located in the NPC basket region oriented with their narrow ends toward the NPC density. Both capsids display a dense interior. (C′) Same as in (C, inset i) but displayed segmented and isosurface rendered.

Figure 3A depicts CLEM of a strongly fluorescent cluster directly associated with the nuclear border of an infected MDM. A slice through the tomographic reconstruction at the correlated position is shown in Figure 3A’’. The segmented and isosurface-rendered representation of this region (Figure 3A’’’) features a total of four cone-shaped and visually intact HIV-1 capsids completely covered in the resin section at different stages of nuclear import (see inserts in Figure 3A’’ and supplementary movie 2). All four capsids that had entered into the NPC were oriented with their narrow end first. We sometimes detected capsids positioning their broad end towards the nuclear envelope in the vicinity of nuclear pores (Figure 3A’’’ left), but this orientation was not observed for particles that had started penetrating the pore.

Capsid-like structures deep inside the central channel (Figure 3A) and on the nucleoplasmic side (Figure 3B, C) retained apparently normal cone-shaped morphology (except for one tubular structure) and dense material inside without any obvious defects (Figure S1). All 32 capsids inside the central channel were oriented with their narrow end towards the nuclear side (Figure S1). Contrary to expectation, capsids on the nucleoplasmic side were found to be oriented in two orientations at equal frequency: either with their narrow end away from (Figure 3B, inset ii) or towards the NPC (Figure 3C) (Figure S1). This result indicated that after passage through the central NPC channel, the HIV-1 capsid remains flexibly attached to the nuclear basket for some time before being released into the nucleoplasm and further trafficking to the site of integration.

### The cyclophilin A binding site is occupied on cytosolic capsids

To investigate the structure of the capsid and its interaction with nuclear pores in more detail, we used cryo electron tomography (cryo-ET). We subjected MDM infected with NNHIV for 48 h and control uninfected cells to specimen thinning by focused ion beam (FIB) milling and acquired 93 and 52 tilt series, respectively (Figure S2, Table S1). In the reconstructed tomograms, we identified 13 enveloped particles outside of cells and 36 capsids inside of cells, either in the cytosol, the nucleoplasm or associated with the NPC (Figure 4, see also Video S3); ten of these capsids were located in direct proximity or inside of nuclear pores (Figure 5A). The capsids appeared to be morphologically intact, and the large majority displayed interior density as typically observed for the nucleoprotein complex.

**Figure 4.**
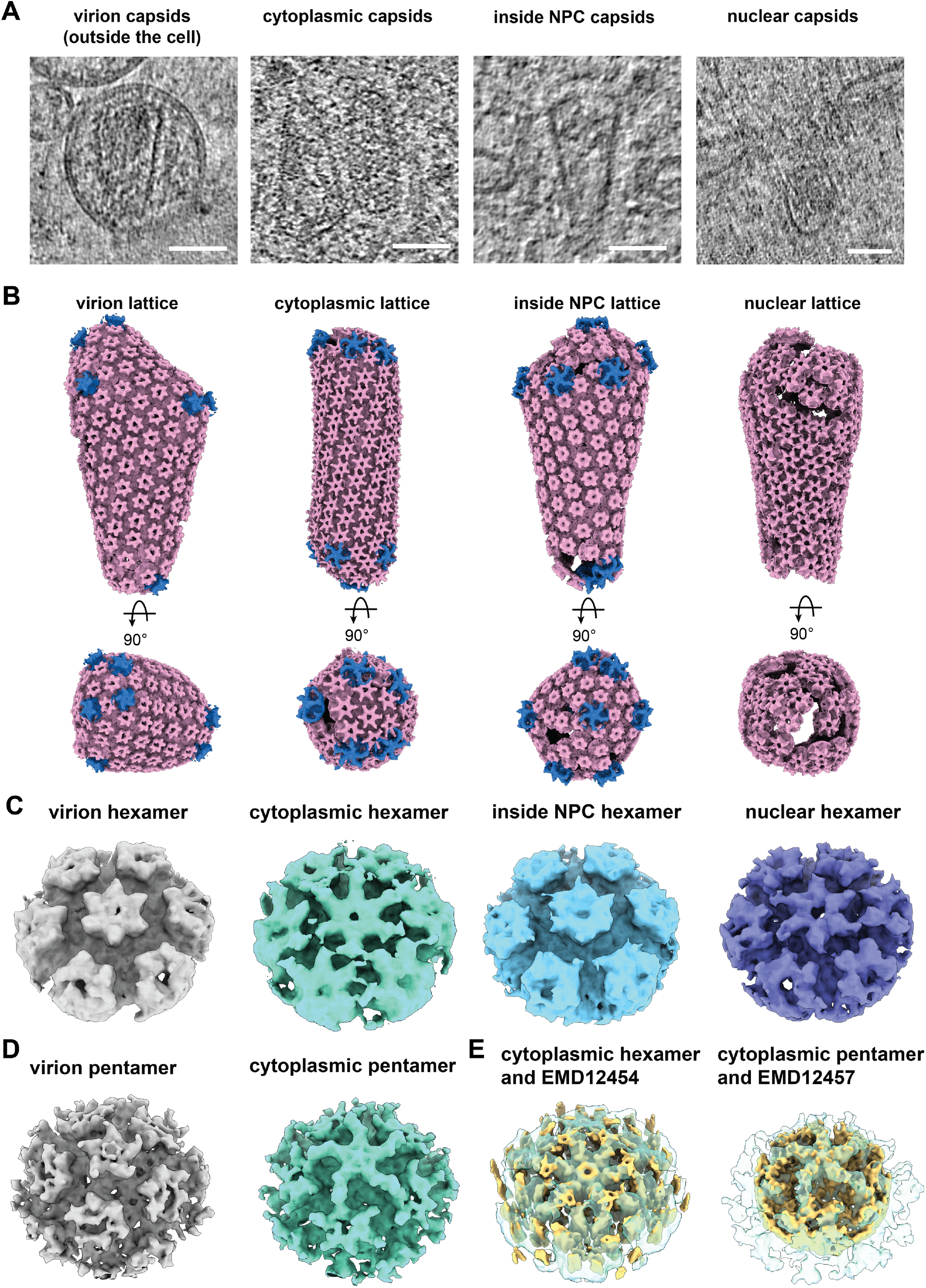
CA lattice is detected in the cytoplasm, during and post nuclear entry. (A) Exemplary virtual slices through cryo-tomograms of HIV-1 capsids inside virions, in the cytoplasm, inside NPCs and in the nucleus. (B) Hexameric capsid lattice traced by STA of CA hexamers in virions, the cytoplasm and inside the central channel and the nucleoplasm. Exemplifying capsids are displayed in side view and a top view looking at the wide end of the capsid. Respective subtomogram averages are shown in pink for the hexamers and in blue for the pentamers. The positions of pentamers were inferred from the hexagonal lattice, where possible, and subsequently subjected to STA. For capsids inside of NPCs, the number of detectable pentamers was insufficient for independent STA. Instead pentamers were combined with those from cytoplasmic capsids. (C) STA of the HIV-1 CA hexamer for virion (grey), cytoplasmic (mint), inside NPC (light blue) and nuclear capsids (purple) shows additional density between the individual CA hexamers only for the cytoplasmic average. (D) STA of the HIV-1 CA pentamer for virion (grey) and cytoplasmic (mint), shows additional density between the central CA pentamer and its neighbors for the cytoplasmic average. (E) Published in vitro structures of CA hexamer and pentamer with CypA bound (EMD12454 and EMD12457 respectively ^40^, both yellow) are shown overlayed with the cytoplasmic capsid averages from this study (both mint). The additional density between neighboring subunits in the cytoplasmic capsids matches where CypA binds in the in vitro structure. Scale bar in (A): 50 nm.

**Figure 5.**
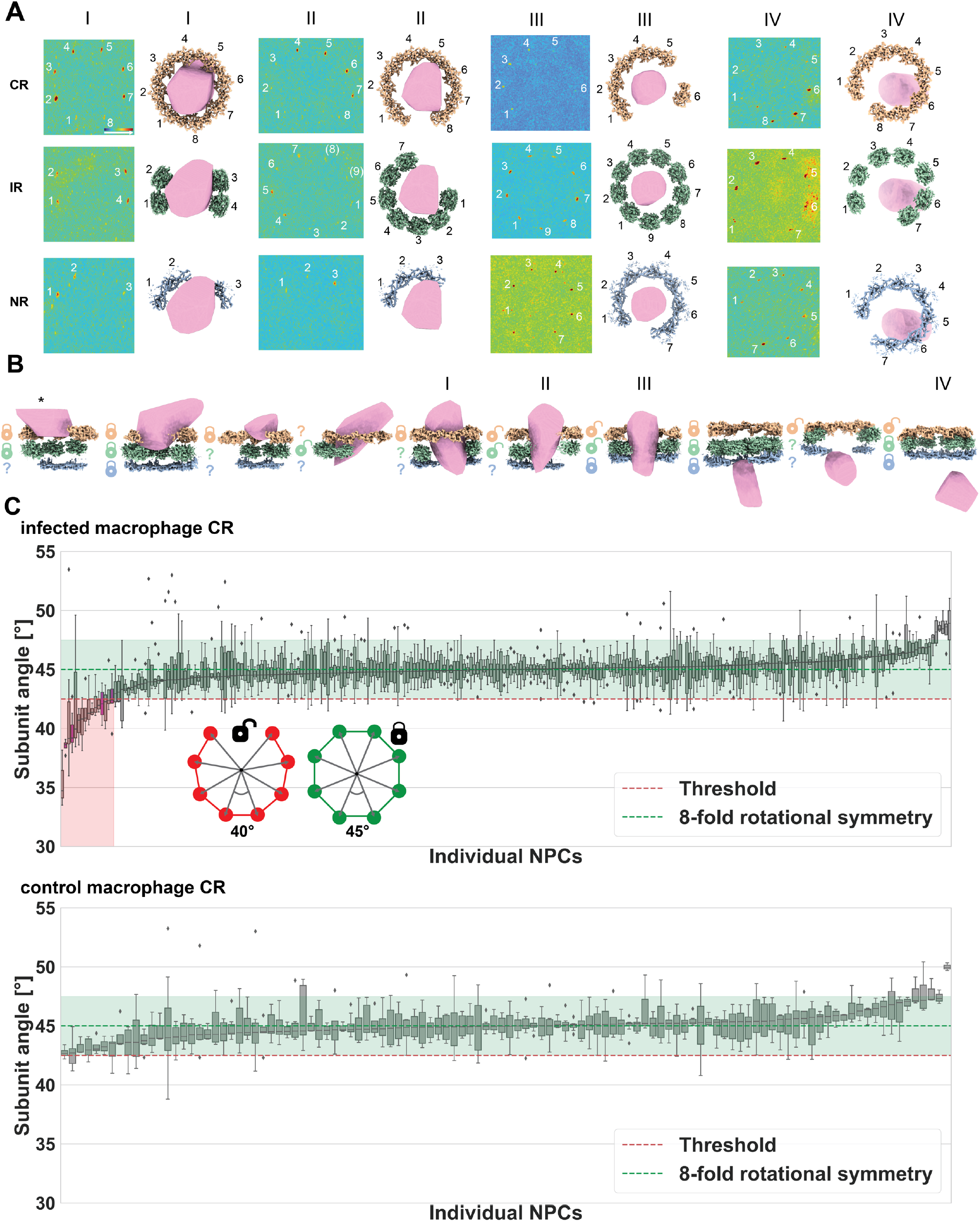
The presence of HIV capsid inside the NPC is linked to abnormal NPC subunit arrangements. (A) Template matching results for subunits of all three NPC rings (CR (light orange, top row), IR (green, middle row), NR (blue, bottom row) for four exemplary NPCs that are associated with an HIV capsid (surface rendering, pink). For each NPC a maximum intensity projection of the constrained cross-correlation (CCC) volume with peaks for that ring subunit is shown color-coded (low CCC values in blue and high values in red). The respective subunit arrangements are indicated in the right panels. (B) Cross section gallery of NPCs sorted according to capsid position. The locks next to each NPC ring represent the status of that ring as either being unperturbed (closed lock) or cracked, the latter meaning inconsistent with the canonical 8-fold rotational symmetry (open lock). This assignment is based on the IAOP (see methods) and a threshold of 42.5°. The asterisk denotes a case where the capsid is only partially contained in the tomographic volume. Question marks represent cases where the number of detected subunits was insufficient for assignment (less than three angle measurements) into either category. For the infected MDM dataset, NPCs without associated capsid were C8-symmetric 247 times and had cracked CR rings twelve times. NPCs with associated capsid were significantly more often cracked (Fisher’s exact test, p=0.0009; four out of nine cracked CR rings). (C) Comparison of the median IAOP per NPC for the CR of HIV-infected and control uninfected MDM. The angles for each CR are plotted as boxplots and sorted by median and are shown in the red transparent box if below a threshold of 42.5° (purple boxes had capsid associated). Angles consistent with C8-symmetric rings are shown in the green transparent box (42.5° - 47.5°). This signature was significantly more frequent in CRs of infected as compared to control MDMs (Fisher’s exact test, p=0.0043; 252 C8-symmetric and 16 cracked rings in infected MDM and 112 C8-symmetric and zero cracked rings in control MDM).

We used subtomogram averaging (STA) of capsid surfaces as previously described ^15, 39^ to structurally analyze the capsid lattice *in situ*. This analysis identified to a large extent the expected hexagonal signature of the CA lattice as well as CA pentamers inside virions, in the cytosol and also for capsids inside the central channel of the NPC (Figure 4B). The lattice was clearly detectable in nuclear capsids as well, but it appeared overall less complete (Figure 4B). Whether these local differences in the nucleoplasm are caused by reduced contrast due to molecular crowding, or alternatively by partial loss of capsid integrity cannot be judged at present.

We hypothesized that the CA lattice might be decorated by different binders during the infection process and categorized our subtomograms according to the subcellular localization of the capsid. As expected, the structure of the CA hexamer and pentamer in enveloped particles outside of cells was very similar to previously analyzed purified particles ^15^, (Figure 4C, D). Inside the cytosol, hexamers and pentamers were also clearly detected but their structure contained additional density consistent with published in vitro structures of CypA bound to CA ^11,40^, in both hexamers and pentamers (Figure 4C,D,E). The respective density was strongly reduced in subtomogram averages obtained from capsids inside the central NPC channel and in the nucleoplasm (Figure 4C), supporting a model in which CypA is bound to the majority of CA hexamers in the cytosol and stripped from the capsid upon NPC entry.

### The NPC scaffold in macrophages is wider than most capsids

Previous analysis in the SupT1 T-cell line by STA pointed to an overall dilated (64 nm at the IR) ^23^, but otherwise normal three-ringed NPC architecture both in infected and control cells, and the diameter of NPCs engaged with capsid was within the overall observed range ^23^. However, the number of capsids observed inside of NPC by cryo-ET was small and these capsids contained the A77V mutation defective in CPSF6 binding. It has been argued that capsids may have to induce additional NPC widening in order to pass through the central channel ^41^. To analyze NPC architecture and width of the central channel in MDM, we subjected 200 and 118 NPCs from infected and control cells, respectively, to STA. The resulting overall structure was reminiscent of previous analyses in Hek293 and SupT1 cells ^23, 26^ (Figure S3A, B). No obvious structural differences were apparent between NPCs from infected and control cells, nor was their NPC diameter significantly different at the IR (Figure S3B, C). However, NPCs in MDM were on average wider than in the previously analyzed SupT1 cells ^23^ (Figure S3B). The IR displayed a more variable diameter as compared to the CR and NR, consistent with the fact that the IR spokes are flexibly connected ^26^. At the given inner diameter of the scaffold in macrophages (∼65 nm, Figure S3D) the NPC would appear to be wide enough to accommodate most HIV-1 capsids with an average diameter of ∼60 nm at their wide end (Figure S2F). To which extent the bulk of FG-Nups contributes to the effective diameter could not be estimated from this analysis however, because it emphasizes scaffold Nups, while FG-Nups are averaged out.

### Nuclear pores crack upon passage of the HIV-1 capsid

Recent advances in template matching technology have allowed us to identify subunits of individual nuclear pores ^42, 43^. To structurally analyze the scaffold of those NPCs that are engaged with a capsid, we used rotational segments of the CR, IR and NR for template matching, as previously described ^43^. Visual inspection of the constrained cross-correlation (CCC) volumes revealed that five out of the ten NPCs engaged with capsid had a distorted scaffold, in which the subunit positioning was not entirely in agreement with the geometry of the canonical 8-fold rotational symmetry (Figure 5A, B). These distortions did not occur homogeneously distributed across the ring structure, but rather at a specific position where the interior angle of the polygon (IAOP) (cartoon in Figure 5C), i.e. the rotation around the symmetry axis from a given to the neighboring subunit, was increased. Concomitantly, this angle was compressed for remaining subunits, thus indicating a rupture event that had cracked the ring structure of the scaffold. Such NPC cracking events were not observed in the three cases where capsids were found associated with the cytosolic face of the NPC, but only in NPCs in which the capsid had penetrated considerably into the central channel. In two NPCs with a capsid deep in the central channel, template matching indicated 9 subunits of the IR (Figurer 5A, II and III, Figure S3E, Video S4). More frequently, however, ring cracking events co-occurred with apparent subunit losses. These were also observed in NPCs that showed a capsid associated with the nuclear face, indicative of damage induced during nuclear entry of this capsid.

### NPC cracking is specific to infected macrophages

Nuclear entry of HIV-1 is a rare event and we wanted to carefully assess the above-described observations for statistical significance. We therefore turned to an objective metric to quantify the number of cracking events in each of the three rings of the NPC. We defined NPC ring cracking events based on the compression of the IAOP of the majority of subunits of a given NPC (Figure 5C, see above). Using this sensitive assay, we first asked whether NPCs in infected MDM that had an associated HIV-1 capsid were more likely to display ring cracking compared to NPCs not associated with a capsid from the same cells. This was clearly the case (Fisher’s exact test, n=268, p=0.0009 for the CR, see legend of Figure 5 for values). We furthermore showed that cracking occurred more frequently in NPCs in infected as compared to control cells, regardless of whether a capsid was found close by the respective nuclear pore (Figure 5C) (Fisher’s exact test, n=380, p=0.0043 for the CR, see legend of Figure 5 for values). At the chosen angular threshold, no cracking events of the CR were observed in non-infected cells (Figure 5C), and they were rare at the level of the IR and NR in these cells (Figure S4).

These observations support the notion that passage of HIV-1 capsids induces NPC cracking. An alternative hypothesis would be that exceptionally large capsids are trapped at NPCs and induce damage, while normal-sized capsids would progress faster into the nucleus and thus were invisible to our analysis. We therefore analyzed whether the size of capsids associated with nuclear pores exceeded that observed for enveloped particles outside of cells. This was not the case (Figure S2F). The notion that nuclear entry of capsids is a rate limiting step in MDM was further supported by our light microscopic and CLEM data presented above; these experiments identified a high number of capsids at different positions of nuclear pores, including the nucleoplasmic side, and showed that these capsids were visually intact and of similar size as observed for intra-virion capsids.

### HIV capsids clash with NPC scaffolds in molecular dynamics simulations

As the capsid approaches the NPC, each CA hexamer can be bound by one of the ∼6,000 FG-repeats. However, the balance between the resulting force pulling the capsid into the central channel and the counterforces from displacing the bulk FG-Nups and clashes with the NPC scaffold remains unclear. Molecular dynamics (MD) simulations in combination with FRET measurements of FG-Nups have recently shed light on their conformational dynamics *in situ* ^44^. Here, we adapted this simulation framework to analyze the steric requirements for nuclear entry of HIV-1 capsids, the response of the FG-Nup network, and the associated forces.

We performed coarse-grained MD simulations to gain a detailed view on the passage of HIV-1 capsids through intact, dilated, and cracked NPCs with and without FG-Nups. We first built an atomic model of a cone-shaped HIV-1 capsid of typical size by completing its well-resolved fullerene-like lattice of CA hexamers and pentamers (Figure S5A,B). We then built a model of the dilated (in-cell) human NPC including FG-Nups (Figure 6A) as described previously for the constricted conformation ^26, 44^. The interaction of capsid and FG-Nups was matched to experiments ^17, 44^ (Figure S5C-E). To model NPC cracking, we split the scaffold and widened it (Figures 6A-C and S7A-C). To cover the NPC diameter increase seen in MDM (Figures S3B), we constructed intact, dilated scaffolds (Figure S7D-F).

**Figure 6.**
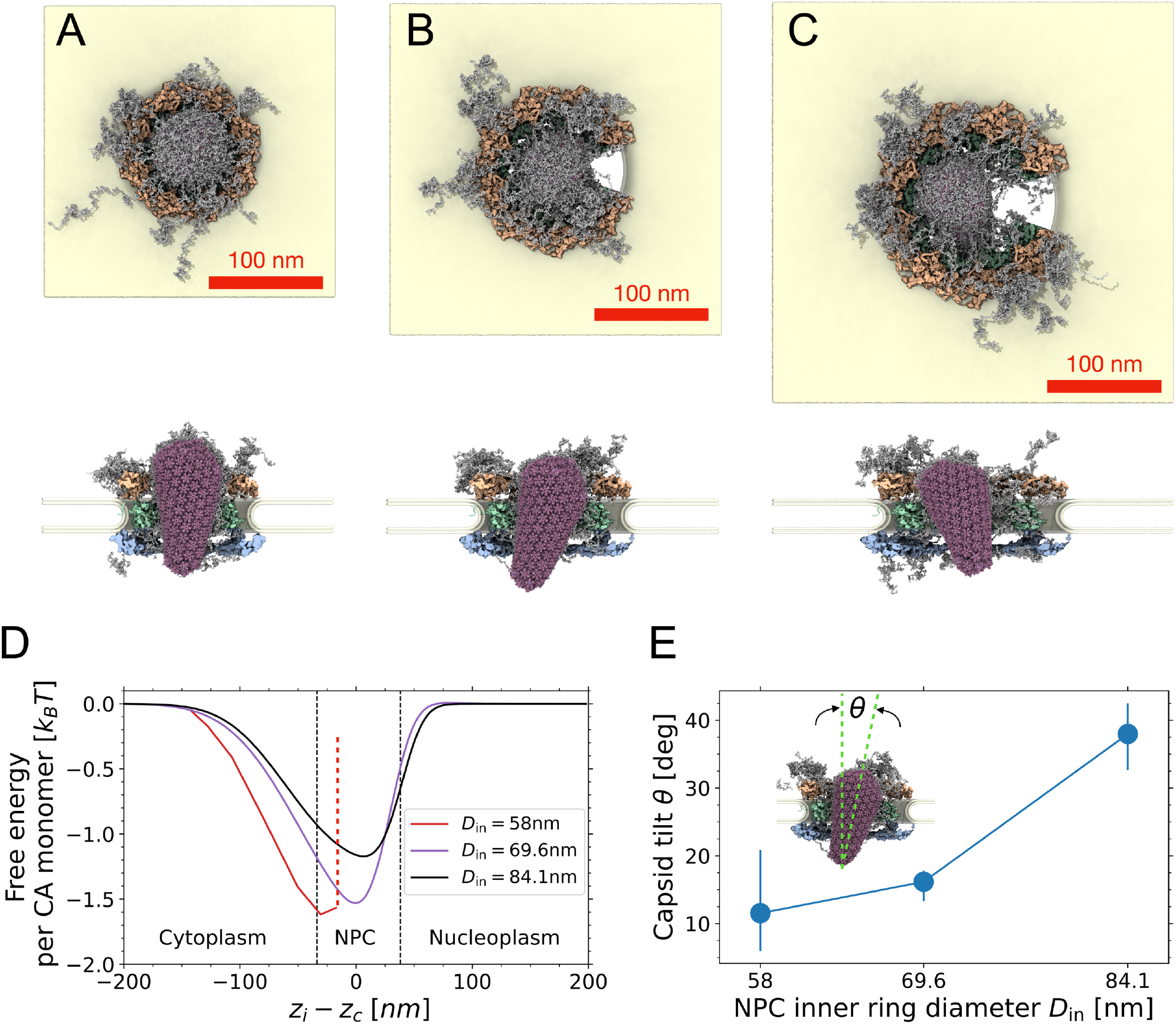
NPC cracking facilitates HIV capsid passage. (A-C) Snapshots of MD simulations of HIV capsid (violet) in intact NPC (A) with inner ring diameter *D*_in_ = 58 nm, and cracked NPCs with *D*_in_ = 69.6 nm (B) and *D*_in_ = 84.1 nm (C). Views from the cytosol (top) and the side (bottom) show end states of the simulations (FG-Nups: grey; CR: orange; IR: green; NR: blue; nuclear envelope: yellow). (D) Free energy profiles for HIV capsid passage through the NPC as a function of the vertical distance *z*_*c*_ − *z*_*i*_ between the capsid center, half-way between its narrow and wide ends, and the inner ring (see Figures S7A-C for the capsid positions used to calculate the mean force shown in Figure S7D). Black dashed lines mark the extent of the NPC, and the vertical red dashed line indicates the position of the steric blockage of the HIV capsid inside the intact NPC. (E) Tilt angle (inset) of capsids released inside intact and cracked NPCs. Symbols and error bars indicate the average, minimum and maximum at the end of three replicas. The interaction strengths 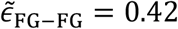 and 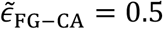 between FG-Nups and CA hexamers and pentamers were matched to experiments. The snapshots were rendered using VMD ^45^.

With these different NPC scaffold models, we identified the minimal steric requirements for HIV-1 capsid passage. In MD simulations of NPCs lacking the FG-Nup network (Figure S6G-L), we found that steric clashes prevent passage of the capsid through the intact in-cell NPC scaffold ^26^. Passage required either dilating the IR diameter further to about 70 nm (Figure S6J), as seen in ∼50% of MDM NPCs (Figure S3D), or cracking the NPC scaffold (Figure S6K,L). The additional mass of FG-Nups is expected to amplify this effect.

Therefore, for a more realistic description of capsid passage, we included the FG-Nup network with interactions matched to experiments. In our MD simulations, FG-Nups in intact and cracked NPCs readily latched on to the HIV-1 capsid, effectively increasing the capsid diameter (Figure 6A-C, see also Video S5). Force-driven simulations (Figure S6G-L) and free energy profiles for capsid passage (Figures 6D and S7) confirmed that capsid passage is possible in the cracked and dilated states with diameters ≥70 nm, but sterically blocked in the ∼58-nm wide in-cell NPC. However, favorable interactions with the FG-Nups resulted in distinct free energy minima for the bound state (Figure 6D), and require further widening for the release of capsid (see Figure S6J-L The favorable interactions with the FG-Nups caused the immersed capsid to tilt sideways (Figure 6E). ^23^

From the MD simulations and consistent with earlier modeling ^41^, we conclude that capsids face two distinct and substantial energetic barriers to exit from the central channel into the nucleus, one caused by repulsive steric interactions with the NPC and the other by the attractive interactions of the capsid with FG-Nups. As found here, the steric barrier can be relieved by NPC cracks, whereas the attractions could be broken either by capsid rupture and release of the HIV-1 genetic material into the nucleus or, alternatively, by competitive binding of additional factors such as Nup153 and CPSF6 to the CA lattice.

## Discussion

The data presented here allow us to assess the different scenarios for the nuclear entry of HIV-1 capsids introduced above. (i) By STA we did not identify deformation of the capsid lattice within the central channel of the NPC as compared to capsids localized elsewhere. (ii) We also did not observe breakage and morphological alteration of the capsid lattice inside the central channel. Instead, we obtained compelling evidence for scenario (iii): Passage of the HIV-1 capsid through the NPC channel of MDM alters the NPC scaffold. Our findings support a model (Figure 7) in which capsids decorated with CypA dock to the cytoplasmic face of the NPC, where CypA is stripped off, most likely by competitive interaction with the Cyp domain of Nup358 in the CR. The capsid is then drawn into the central channel by saturating the respective FG-binding sites towards the broad end of the capsid ^31^ As it penetrates more deeply, clashes with both the NPC scaffold and FG-Nups emerge. Entry into the limited space creates lateral force that in turn can lead to stretching of the NPC scaffold. NPC stretching may then extend the ring structure to a degree that it cracks, likely allowing a spatial re-distribution of FG-Nups and further progression of the capsid through the nuclear pore. We cannot exclude that capsid elasticity of scenario (i) could occur transiently and may thus contribute to nuclear entry of the capsid as well. Furthermore, potential capsid breakage as in (ii) could lead to complete capsid disintegration, thereby escaping detection. However, based on the high proportion of cracked pores and the observation of morphologically intact capsids in the nucleus, we propose that HIV-1 capsids pass the NPC more or less intact and, if sterically blocked, crack its rings.

**Figure 7.**
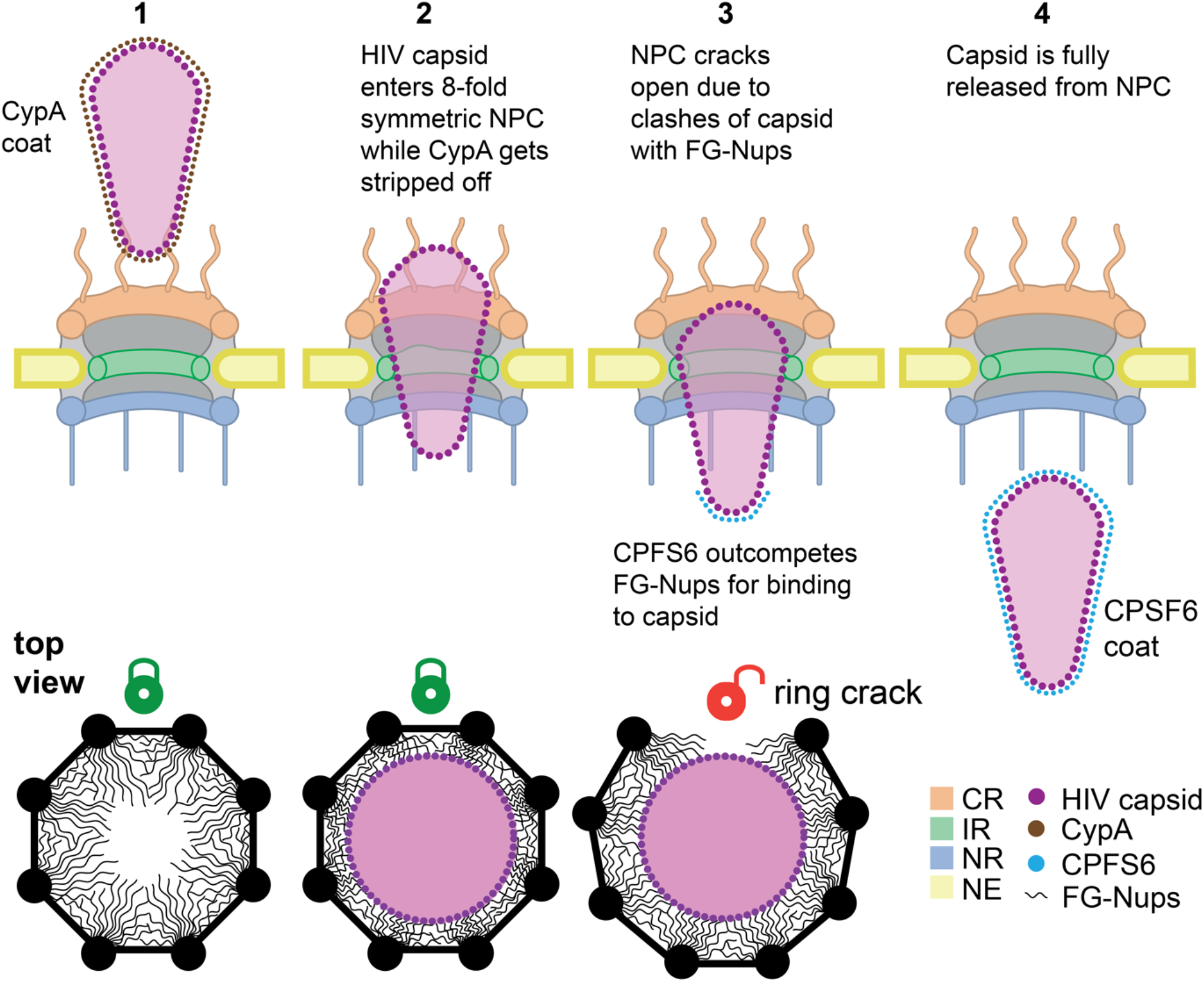
Model for nuclear entry of the HIV-1 capsid in macrophages. (1) Side view cross section: A cone-shaped HIV-1 capsid decorated with CypA is oriented with its narrow end towards the cytoplasmic face of the NPC. The cytoplasmic filaments of the CR guide and orient the capsid for further entry into the pore. The top view cross section below shows the intact (green closed lock) C8-symmetric ring structure with the FG-Nups extending into the central channel. (2) Side view cross section: The cone-shaped capsid is partially inserted into the central channel of the NPC, thereby losing the associated CypA. Its wide end has not yet passed through the narrowest point of the NPC scaffold (CR opening). The top view shows the intact (green closed lock) C8-symmetric ring structure with the capsid inserted. The FG-Nup mesh network is compressed due to the capsid insertion. (3) Side view cross section: The capsid has passed through the CR opening by cracking the NPC scaffold and is now fully inserted into the widened central channel of the NPC. CPFS6 starts to coat the tip of the capsid as it reaches the nucleoplasm by outcompeting FG-Nups. The top view shows the cracked (red open lock) ring structure with the capsid inserted. The FG-Nup mesh network relaxes due to the ring crack. (4) Side view cross section: The capsid has been released from the nucleoplasmic side of the NPC and has acquired a full CPFS6 coat. The HIV-1 capsid, CPFS6 and the individual rings and FG-Nups of the NPC are shown color-coded.

We speculate that eventual capsid progression towards the nucleoplasm may be driven by interactions with Nup153 FG-repeats. At the nuclear basket, the capsid may transiently remain latched to the very long Nup153 FG-repeats and thus rotate in the chromatin-free region, as occasionally observed in the simulations. This would agree with in vitro measurements of binding constants of different FG-Nups ^17, 46^, and can explain both the observed delayed release into the nucleoplasm and the variable orientation of nuclear basket associated capsids. Ultimately, FG binding sites in the capsid lattice will be competitively saturated by abundant nuclear CPSF6, causing capsid release from the nuclear basket to migrate deeper into the nucleoplasm. This model is consistent with CPSF6 gene silencing experiments and mutation of the CPSF6 binding site in CA; in these cases, capsids were prominently observed close to the nuclear basket region and differences in HIV-1 integration sites were found ^20, 23^.

We had previously found that cone-shaped capsids enter dilated NPCs in T-cells ^23^. However, cryo-ET had captured only few events, and thesecapsids contained the A77V mutation defective in CPSF6 binding. Moreover, by analyzing the NPC architecture through subtomogram averaging, differences across a small set of individual particles could not be readily detected. Here, by using template matching of NPC subunits ^42,43^ we could reliably detect damage of individual NPCs and, by greatly expanding the statistics, connect NPC cracking to HIV capsid passage.

That the HIV-1 capsid cracks the NPC was not necessarily intuitive, because the NPC scaffold is an elaborate structure designed to persist strain laterally imposed by the nuclear membranes ^25,43,47^, while the capsid has to disintegrate at some stage during the post-entry process. The exact forces remain challenging to measure experimentally. In the MD simulations, forces of 10-100 pN pull the capsid into the NPC central channel (Figure S7D), which is in the range of forces required for sub-second unfolding of proteins ^48^. The cone shape of stuck capsids converts this axial force into a radial force that could drive NPC cracking. Instead of the NPC cracking the capsid, the capsid cracks the NPC.

Deviations from the canonical 8-fold rotational symmetry of the NPC occur only rarely under native conditions ^49^. A recent study investigating NPC subunit arrangement and diameter during stem cell differentiation has detected abnormal rotational symmetries and NPC overstretching as a result of knock out of the scaffolding Nup133 ^43^. In contrast to this study, however, straining of the NPC scaffold by the HIV-1 capsid occurs from the inside. The observed distortions are markedly different, indicating that the changes observed by Taniguchi et al. are of different origin. Specifically, Taniguchi et al. did not observe rotational symmetry mismatches across the three rings within individual NPCs, but rather NPCs were overstretched and disintegrated, thereby detaching from the nuclear membranes ^43^. Ring cracking by HIV-1 capsid passage shows specific features: subsequent to cracking of one ring, the scaffold contacts with the next ring do not rotationally register. This finding is intriguing because it emphasizes that the NR and CR have intrinsic stability. Furthermore, our results show that nuclear pores may transport cargos that are larger than their diameter.

The findings presented here provide an explanation for the evolution of the unique HIV-1 capsid structure. Protecting the genome and allowing reverse transcription inside a protective shell requires a closed structure, but not this cone-shaped geometry. However, the complete capsid also acts as the importin for nuclear entry of the HIV-1 subviral complex ^32, 33^, and this is absolutely essential for infection of target cells that do not break down their nuclear envelope during mitosis. Shape most likely reflects this function: the narrow end facilitates threading into the narrow channel of the NPC, and the increasing width of the cone allows to accommodate the volume requirement for the genome and provides avidity effects by an increasing number of FG binding sites on the capsid lattice as it progresses into the NPC. Maximum width may then be a trade-off between what is absolutely needed to package the genome (most likely together with partial cDNA due to cytoplasmic reverse transcription) and the size limit to pass through the (partly disintegrated) central NPC channel. Thereby, NPC cracking might be a simple byproduct of HIV-1 nuclear entry with optimal payload, without any direct consequences for infectivity. Importantly, damage to individual NPCs must be compatible with cell viability as infected macrophages can survive for prolonged time. Could there be a further benefit from NPC cracking for the virus? One may speculate that the capsid lattice becomes mechanically prepared for rupture while it sheers at the NPC scaffold, but does not immediately disintegrate because of rapid surface coating with CPSF6 upon reaching the nuclear side. Whether this is the case, and whether there may be other benefits for the virus, e.g. inducing nuclear envelope rupture and thus locally changing the nuclear proteome, remains to be further investigated.

## Materials and Methods

### Cells

Human embryonic kidney 293T cells (HEK293T) ^50^ were maintained in Dulbecco’s modified Eagle medium (DMEM; Thermo Fisher Scientific) supplemented with 10% fetal bovine serum (FBS; Capricorn Scientific GmbH, Germany), 100 U/ml penicillin and 100 mg/ml streptomycin (Thermo Fisher Scientific). Monocyte-derived macrophages (MDM) were obtained from human peripheral blood mononuclear cells (PBMC) isolated from buffy coats of healthy donors as described previously ^20^. Buffy coats were obtained from anonymous blood donors at the Heidelberg University Hospital Blood Bank according to the regulations of the local ethics committee. MDM were cultivated in RPMI 1640 medium (Thermo Fisher Scientific) supplemented with 10% heat inactivated FBS, antibiotics (as above), and 5% human AB serum (Capricorn Scientific GmbH, Germany). Cells were cultivated at 37ºC in a humidified incubator with a 5% CO2 atmosphere. For seeding, MDM were detached by Accutase (StemCell Technologies) according to the manufacturer’s instructions.

### Plasmids

Plasmid pNNHIV for production of non-infectious, reverse transcription competent HIV-1 particles ^23^, the proviral plasmids pNL4-3 ^51^ and pNL4-3ΔEnv ^20^, and the Vpr.IN_D64N/D116N_.mScarlet fusion protein encoding plasmid pVpr.IN_D64N/D116N_.mScarlet ^23^ were described previously. Plasmid pEnv-4059 encoding an R5-tropic Env from a clinical HIV-1 isolate ^52^, was kindly provided by R. Swanstrom (University of North Carolina, Chapel Hill, NC, USA). Plasmid pVpr.IN.eGFP encoding a Vpr.IN.eGFP fusion protein with an HIV-1 protease recognition site between Vpr and IN ^53^, was kindly provided by A. Cereseto (CIBIO, Mattareo, Italy). Plasmid pNNHIVΔEnv contains a 1 bp fill-in of an StuI site in the *env* ORF resulting in a frameshift and premature stop codon (primers used for PCR: forward, 5’-CAGACAGGCCTCGTCCAAAGGTATCCTTTGAG-3’; reverse, 5’-CAGACGCTAGCTATCTGTTTTAAAGTGGCATTC-3’).

### Antibodies

For immunofluorescence staining, rabbit polyclonal antiserum against HIV-1 CA raised against purified recombinant protein (in house) ^54^, mouse monoclonal antibody against human lamin A/C (sc-7292; Santa Cruz), mouse monoclonal antibody against Nup153 (ab24700; Abcam) and mouse monoclonal antibody against FG-Nups (Mab414) (ab24609; Abcam) were used at a dilution of 1:1,000, 1:100 and 1:200, respectively. For spinning disc confocal microscopy, secondary antibodies donkey anti-rabbit IgG and donkey anti-mouse IgG conjugated with Alexa Fluor 488 and 647 (Thermo Fisher Scientific), respectively, were used at 1:1,000 dilution. For STED microscopy, secondary antibodies goat anti-rabbit IgG and goat anti-mouse IgG conjugated with Atto 594 (Merck; 77671) and Abberior® STAR RED (Merck; 52283), respectively, were used at 1:200 dilution.

### Virus and virus-like particles

HIV-1 virions and non-infectious virus-like particles were produced in HEK293T cells grown on 175 cm^2^ tissue culture flasks (side bottom) transfected with respective plasmids (total 70 μg DNA per flask) using calcium phosphate transfection. For production of infectious R5-tropic HIV-1, cells were transfected with pNL4-3ΔEnv and pEnv-4059 at a molar ratio of 4.5:1. To produce IN.eGFP labeled R5-tropic HIV-1, cells were transfected with pNL4-3ΔEnv, pVpr.IN.eGFP and pEnv-4059 at a molar ratio of 4.5:1:1. For production of non-infectious IN.mScarlet labeled R5-tropic NNHIV, cells were transfected with pNNHIVΔEnv, pVpr.IN_NN_.mScarlet and pEnv-4059 at a molar ratio of 4.5:1:1. Medium was changed 6 h after transfection and cells were further incubated at 37ºC and 5% CO_2_. At 44 h post-transfection, culture media from virus-producing cells were harvested and cleared by filtration through a 0.45 μm nitrocellulose filter (Carl Roth, Germany), and particles from media were concentrated by ultracentrifugation through a 20% (w/w) sucrose cushion for 90 min at 27,000 rpm (at 4ºC) in a Beckman SW32 rotor (Beckman Coulter Life Sciences). Particles were resuspended either in PBS or in PBS containing 10% FBS and 10 mM HEPES (pH 7.2), then aliquoted and stored at –80ºC. Particle-associated RT activity was determined by SYBR Green-based Product-Enhanced Reverse Transcription assay (SG-PERT) ^55^.

### Sample preparation for confocal and STED microscopy

MDM seeded in 8-well μ-Slides with a glass bottom (ibidi GmbH, Germany) were infected with non-labeled or fluorescently labeled wild-type HIV-1 or NNHIV (5–15 μUnits of RT/cell) pseudotyped with R5-tropic 4059 Env and incubated at 37°C until indicated time post-infection. For both spinning disc confocal and super-resolution STED microscopy, samples were fixed with 4% FA in PBS (15 min), rinsed with PBS, permeabilized with 0.5% Triton X-100 in PBS (5–20 min) and washed three times (5–10 min) with PBS. For detection of nuclear HIV-1 CA, cells were further extracted using ice-cold methanol for 10 min (at –20ºC) and subsequently washed twice with 3% bovine serum albumin (BSA) in PBS. Samples were blocked with 3% BSA in PBS for 30 min and staining with primary and secondary antibodies (both diluted in 0.5% BSA in PBS) was carried out at room temperature for 1 h each. When relevant, cell nuclei were visualized by DNA staining with 0.2 μg/ml Hoechst33258 (Thermo Fisher Scientific) in PBS for 30 min.

### Spinning disc confocal microscopy

Spinning disc confocal microscopy (SDCM) was performed using a Nikon Ti Perkin Elmer Ultra VIEW VoX 3D spinning disk confocal microscope (Perkin Elmer, MA, USA) equipped with a 100× oil immersion objective (NA 1.4; Nikon). Multichannel z series of randomly selected cells were acquired with a z-spacing of 200 nm and excitation with the 405-, 488-, 561-, and 640-nm laser lines, using the Volocity software (Perkin Elmer, MA, USA).

### STED microscopy

Stimulated emission depletion (STED) imaging was performed with a 775 nm Expert Line STED system (Abberior Instruments GmbH, Germany) equipped with an SLM-based easy3D module and an Olympus IX83 microscope using a 100× oil immersion objective (NA 1.4; Olympus UPlanSApo). Dual-color STED images were acquired line sequentially, using the 590- and 640-nm excitation laser lines and two APD spectral detectors set to collect photons with the wavelength between 590–630 nm and 650–720 nm, respectively. Acquisitions using 405- and 488-nm laser lines were in confocal mode only. Nominal STED laser power was set to 15–40% of the maximal power of 3 W with 10 μs pixel dwell time, 15 nm pixel size and 9× accumulation. Acquired STED images were deconvoluted with Huygens Deconvolution software (Scientific Volume Imaging) using Classic Maximum Likelihood Estimation (CMLE) algorithm and Deconvolution Express mode with ‘Conservative’ settings. For 3D STED data acquisition, 30% of the STED laser power was used for fluorescence depletion in the Z-axis and RESCue illumination scheme was used to minimize bleaching. Sampling frequency was 20 nm in xy axis and 70 nm in z. For sampling of the entire nuclear volume (∼6–10 μm along the optical axis) by 3D STED, 80–150 super-resolved images were acquired. The bleaching during the acquisition was reduced by implementing a light dose management (DyMIN) that specifically activates and modulates the intensity of the STED depletion laser beam to switch off fluorophores only near the fluorescent feature to be recorded ^56^.

### Image analysis

Quantification of CA signals localized at the nuclear envelope or inside the nucleus of infected MDM visualized by SDCM was performed using the Icy software (^57^). The volumes of individual HIV-1 CA signals in acquired z-stacks were automatically detected using the spot detector function. Objects displaying positive signal in the laminA/C or Hoechst channel were classified as nuclear envelope (NE) associated or as intranuclear, respectively. Nuclear signals were further visually examined and manually currated to ensure that objects located in nuclear regions with very low or undetectable Hoechst staining were not excluded from the analysis.

To determine percentage of CA signals colocalizing with FG-Nups at the nuclear perihery in infected MDM visualized by 3D STED, deconvoluted z-stacks were reconstructed using the Imaris software (Bitplane AG, Switzerland). Individual HIV-1 CA signals were automatically detected using the spot detector Imaris module, creating for each distinct fluorescent signal a 3D ellipsoid object with Z axis = 1.5*X,Y axis. For all objects in the proximity of NE, the median signal intensity within objects was quantified for the FG-Nups channel. All objects representing CA signals that had maximum FG-Nups signal intensity higher than 50 (a.u.) were scored as co-localizing. The same threshold was applied to all four datasets. To quantify the total amounts of nuclear pores per nucleus, individual FG-Nups signals were automatically detected by spot detector module as above and counted. To estimate the NPC density, the dimensions of nuclei mid-sections was measured using the Fiji software ^58^ and the surface area of each nucleus was then calculated as the surface of ellipsoid.

### Sample preparation for CLEM

4 × 10^4^ MDM were seeded on carbon-coated sapphire discs (Engineering Office M. Wohlwend, Switzerland) placed in a glass-bottomed ‘microwell’ of 35 mm MatTek dish (MatTek, Ashland, MA, USA) and cultured for 16–24 h at 37ºC. Cells were infected with IN.mScarlet labeled NNHIV particles pseudotyped with R5-tropic 4059 Env at 60 μU RT/cell. At 48 h p.i., infected cells were cryo-immobilized by high pressure freezing using a HPM010 high pressure freezer (BAL-TEC, Balzers, Liechtenstein) and discs with frozen cells were transferred to freeze-substitution medium (0.1% uranyl acetate, 2.3% methanol and 1% H_2_O in Acetone) at –90ºC. Subsequent freeze-substitution and embedding of samples in Lowicryl HM20 resin (Polysciences, Inc., USA) was performed in an EM AFS2 freeze-substitution device (Leica Microsystems) equipped with a EM FSP robotic solution handler (Leica Microsystems) according to Kukulski et al. (^59^), modified as follows: Samples were incubated in FS medium for 5 h at –90ºC and temperature was then raised to –45ºC (at 7.5ºC/h). Samples were washed with acetone (3 × 25 min) and infiltrated with increasing concentrations of Lowicryl HM20 in acetone (25, 50% and 75%; 3 h each), while raising temperature to –25ºC (3.3ºC/h). The acetone-resin mixture was replaced by pure Lowicryl HM20 for 1 h and the resin was exchanged three times (3, 5 and 12 h). Samples were polymerized under UV light for 24 h at –25ºC and polymerization continued for an additional 24 h while the temperature was raised to 20ºC (at 3.7ºC/h).

### CLEM and electron tomography

250-nm thick resin sections were obtained using a EM UC7 ulramicrotome (Leica Microsystems) and placed on a slot (1 × 2 mm) EM copper grids covered with a formvar film (Electron Microscopy Sciences, FF2010-Cu). Grids were placed (section face-down) for 10 min on 20 μL drops of 1 × PHEM buffer (pH 6.9) containing 0.1 μm TetraSpeck beads (1:25) (Thermo Fisher Scientific) serving as fiducial markers and 10 μg/ml Hoechst33258 (Thermo Fisher Scientific) to stain nuclear regions in cell sections. Unbound fiducials were washed off on several drops of water and grids were transferred on 25 mm glass coverslips mounted in a water-filled ring holder for microscopy (Attofluor cell chamber, Thermo Fisher Scientific). Z stacks of sections were acquired with a PerkinElmer UltraVIEW VoX 3D spinning-disc confocal microscope (Perkin Elmer, MA, USA) and then visually examined using the Fiji software ^58^ to identify regions of interest (ROIs). Sections on EM grids were stained with 3% uranyl acetate (in 70% methanol) and lead citrate. Individual grids were placed in a high-tilt holder (Fischione Model 2040) and loaded to a Tecnai TF20 electron microscope (FEI) operated at 200 kV, equipped with a field emission gun and a 4K-by-4K pixel Eagle CCD camera (FEI). To map all cell sections on grid, a full grid map was acquired using the SerialEM software ^60^. To identify ROIs in resin cell sections for image acquisitions, EM images and imported SDCM images were pre-correlated in SerialEM using the fiducials as landmark points ^61^ and single-axis tilt series were acquired in correlated positions using SerialEM with a tilt range from –60° to +60°, angular increment of 1° and a nominal pixel size of 1.13 nm. Alignments and 3D reconstructions of tomograms were done with IMOD software ^62^. High precision post-correlation was performed using eC-CLEM plugin ^63^ in Icy software ^57^.

### Quantitative image analysis of capsids acquired by CLEM-ET

Segmentation, isosurface rendering and quantitative analysis of the capsid interior were done in the Amira software (Thermo Fisher Scientific) as described in Zila et al. ^23^. Briefly, to exclude the CA layer density from the interior of manually segmented capsid, the ‘erosion’ algorithm was used to shrink the volume of segmented structure. The interior voxel intensity median in shrunken volume was then determined using the ‘label analysis’ function and normalized to intensity median of 3–5 volumes placed in the proximity of the structure. Only structures fully covered in the EM section (not truncated at the section edge) were included in the analysis.

### MDM vitrification and cryo-FIB milling

MDM were detached by accutase (StemCell Technologies) according to the manufacturer’s instructions. 4 × 10^4^ MDM were seeded on glow discharged and UV-light sterilized 200-mesh EM gold grids coated with R 2/2 holey carbon films (Quantifoil Micro Tool GmbH), which were placed in a glass-bottomed ‘microwell’ of 35-mm MatTek dish (MatTek, Ashland, MA, USA). After seeding, cells were cultured for an additional 24 h at 37 ºC. For infection, cells were incubated with IN.mScarlet labeled NNHIV particles pseudotyped with R5-tropic 4059 Env at 60 μU RT/cell for 48 hours. Mock-infected and infected MDM were vitrified by plunge freezing into liquid ethane at –183°C using an EM GP2 plunger (Leica Microsystems). The blotting chamber was maintained at 37°C temperature and 90% humidity. Before plunge freezing, 3 μl of culture medium were applied onto grids. Subsequently, the grids were blotted from the back side for 3 s with a Whatman filter paper, Grade 1, and plunge frozen. The samples were then cryo-FIB milled using an Aquilos 2 microscope (Thermo Fisher Scientific) similar to a previously described workflow ^64^. In brief, samples were coated with an organometallic platinum layer using a gas injection system for 10 sec and additionally sputter coated with platinum at 1 kV and 10 mA current for 10 sec. Milling was performed with AutoTEM (version 2.4.2) (Thermo Fisher Scientific) in a stepwise manner with an ion beam of 30 kV while reducing the current from 1000 pA to 50 pA. Final polishing was performed with 30 pA current with a lamellae target thickness of 200 nm.

### MDM cryo-ET data acquisition

Cryo-ET data for infected and control macrophages were collected at 300 kV on a Titan Krios G4 microscope (Thermo Fisher Scientific) equipped with a E-CFEG, Falcon 4 direct electron detector (Thermo Fisher Scientific) operated in counting mode and Selectris X imaging filter (Thermo Fisher Scientific). For each grid, montaged grid overviews were acquired with 205 nm pixel size. Montages of individual lamellae were taken with 30 nm pixel size. Tilt series were acquired using SerialEM (version 4.0.20) ^60^ in low dose mode as 4K x 4K movies of 10 frames each and on-the-fly motion-corrected in SerialEM. The magnification for projection images of 53000x corresponded to a nominal pixel size of 2.414 Å. Tilt series acquisition started from the lamella pretilt of ± 8° and a dose-symmetric acquisition scheme ^65^ with 2° increments grouped by 2 was applied, resulting in 61 projections per tilt series with a constant exposure time and targeted total dose of ∼135 e^-^ per Å^2^. The energy slit width was set to 10 eV and the nominal defocus was varied between −2.0 to −4.0 μm. Dose rate on the detector was targeted to be ∼4-6 e^-^ /px/sec.

### Tomogram reconstruction

The motion corrected tilt series were corrected for dose exposure as previously described ^66^ using a Matlab implementation that was adapted for tomographic tilt series ^67^. Projection images with poor quality were removed after visual inspection. The dose-filtered tilt series were then aligned with patch-tracking in AreTomo (version 1.33) ^68^ and reconstructed as back-projected tomograms with SIRT-like filtering of 15 iterations at a binned pixel size of 9.656 Å (bin4) in IMOD (version 4.11.5) ^62^ From the reconstructed tomograms, those containing nuclear pore complexes and/or HIV capsids were selected by visual inspection.

The same tomograms were also reconstructed with 3D-CTF correction using novaCTF ^69^ with phase-flip correction, astigmatism correction and 30 nm slab. Tomograms were binned 2x, 4x and 8x using Fourier3D [B. Turoňová, turonova/Fourier3D: Fourier3D, Version v1.0, Zenodo (2020)].

### HIV CA hexamer subtomogram averaging

Capsid-like structures were identified in the infected macrophage SIRT-like filtered bin4 tomograms and then manually segmented in napari [napari contributors (2019). napari: a multi-dimensional image viewer for python. doi:10.5281/zenodo.3555620]. The segmentation was used to create a convex hull with the cryoCAT package. Subtomogram averaging of the capsid hexamer was performed similarly to a previously described protocol ^15^. In detail, first the segmented capsid surfaces (at 4x binning) were oversampled with a sampling distance of 2 voxels and each position was assigned an initial orientation that was determined based on the normal vector of the segmented cone surface at given position (the in-plane angle was assigned randomly). Then the coordinates were multiplied by factor of 2 to obtain positions for bin2 tomograms. All subsequent subtomogram alignment and averaging was performed with imposed C6 symmetry in novaSTA [Turoňová, B. turonova/novaSTA: Advanced particle analysis, Version v1.1. Zenodo (2022)].

For virion capsids, the oversampled positions of four capsids were used to extract bin2 subtomograms from 3D-CTF corrected tomograms. These subtomograms were used to generate an initial featureless average and then aligned for six iterations. A distance threshold of 16 voxels was used to remove overlapping subtomograms (i.e. particle duplicates). Low CCC threshold and/or incorrect orientation (both assessed visually in using ArtiaX ^70^ lead to additional removal of subtomograms. The resulting subtomograms were then aligned for six iterations to obtain a better average and lattice arrangement. After further removal of suboptimal positions, the final subtomograms were again aligned for six iterations.

This STA map of the virion hexamer was then used as an existing reference (lowpass filtered to 40 Å) to align all oversampled virion capsid positions and obtain more complete lattices. The resulting positions were used to remove duplicates and suboptimal positions as described above. After multiple iterations of removal of incorrectly oriented positions and then manual addition of potential positions to complete the lattice again in ArtiaX ^70^, the final subtomograms were again aligned for nine iterations (for final particle numbers see Table S1) For capsids in the cytoplasm, inside NPCs or in the nucleus, the same procedure was applied.

### HIV CA pentamer subtomogram averaging

All subsequent subtomogram alignment and averaging was performed with imposed C5 symmetry in novaSTA [Turoňová, B. turonova/novaSTA: Advanced particle analysis, Version v1.1. Zenodo (2022)]. Positions and orientations for subtomogram averaging of the CA pentamer were obtained by placing particles into the areas of the capsid lattice where hexamers were arranged around a pentamer hole as visualized in ArtiaX ^70^. Virion candidate positions were processed separately from cytoplasmic and inside NPC candidate positions. In each case, the candidate positions were used to extract bin2 subtomograms from 3D-CTF corrected tomograms. These subtomograms were used to generate an initial featureless average and then aligned for nine iterations. Low CCC threshold and/or incorrect orientation (both assessed visually in using ArtiaX ^70^} led to additional removal of subtomograms. The final subtomograms were aligned for nine iterations (for final particle numbers see Table S1). For FSC calculations, the virion pentamer was treated as halfmap1 and EMD-3466 ^15^ as halfmap2 and the resolution evaluated at FSC=0.5. The cytoplasmic pentamer map was treated as halfmap1 and EMD-12457 ^40^ as halfmap2 and the resolution evaluated at FSC=0.5.

### HIV CA pentamer template matching

A published CA pentamer STA map (EMD-3466 ^15^) was downsampled to a pixel size of 4.828 Å (bin2 for this dataset) and then used as a search template to perform template matching in parts of a bin2 tomogram with capsid-like structures using GAPSTOP™ [https://gitlab.mpcdf.mpg.de/bturo/gapstop_tm] ^42^. The angular search range was set to 5-degree angular sampling. The highest CCC peaks from template matching were extracted with their corresponding orientations and then visually inspected in ArtiaX ^70^ to obtain only the plausible candidates that are oriented correctly to the capsid surface as visible in the tomogram.

### NPC particle selection and subtomogram averaging

Positions of NPCs were manually selected in the bin4 SIRT-like filtered tomograms as described previously ^71^. Extraction of particles from the novaCTF-corrected tomograms, subtomogram alignment and averaging was performed using novaSTA [Turoňová, B. turonova/novaSTA: Advanced particle analysis, Version v1.1. Zenodo (2022)]. NPC averaging was performed as described previously ^25^. First, an average of the whole NPC was obtained while utilizing C8 symmetry and a cylindrical mask. Next, coordinates of the subunits were extracted based on the aligned positions using C8 symmetry. With these subunit positions an average structure of the asymmetric unit of the NPC was obtained using an elliptical mask. To obtain averages of individual NPC rings, first particle positions were re-centered to the respective area (CR, IR, NR, LR, nuclear basket) based on their position in the subunit average. Then the newly extracted bin4 subtomograms for the individual rings were aligned and averaged with elliptical masks. To generate C8 symmetric composite NPC maps, first the final averages of individual ring maps were fitted into the asymmetric subunit average. Then the composite map was created by applying symmetry based on the coordinates used for splitting the initial whole NPC average into asymmetric units. The entire NPC STA procedure was first performed for control and infected macrophage NPCs separately and then redone for the combined set of NPCs after determining no discernible difference between the two macrophage NPC structures (control and infected).

### NPC diameter measurements

The NPC diameters at different points were measured based on the coordinates obtained from STA maps of individual rings (CR, IR, NR) using previously published MATLAB scripts ^25^. Using ChimeraX ^72^, the measurement point of interest coordinates in the individual ring average were determined with the marker tool and the particle list coordinates were offset by the shift between the center of the average and the measurement point of interest. Only NPCs with five or more subunits were considered for diameter measurement of NPCs. As previously described ^25^, line segments were determined that connected opposing subunits for each individual NPC. The NPC center was defined as the point to which the distance of all line segments was minimal. The NPC radius was calculated as the distance from the center to each subunit. Data was plotted using Prism9 software (Figure S3 C,D). Statistical significance was tested using unpaired two-tailed t test in Prism9.

### NPC ring subunit template matching

The obtained bin4 STA averages for CR, IR and NR were masked to only include a single subunit and then used as search templates (lowpass filtered to 30 Å) to perform template matching in full 3D-CTF-corrected bin4 tomograms with visible NPCs using GAPSTOP™ [https://gitlab.mpcdf.mpg.de/bturo/gapstop_tm] ^42^ based on the STOPGAP software package^73^. The angular search range was set to a 6-degree sampling which has been shown to increase TM performance ^42^. The constrained cross-correlation (CCC) peaks which had values 7x above the mean of the CCC-volume were extracted with their corresponding orientations and then manually cleaned in the ArtiaX plugin ^70^ for ChimeraX ^72^ to obtain only plausible candidates that are oriented correctly in the nuclear envelope. This resulted in three different particle sets for CR, IR, NR with all template matched subunit coordinates and orientations for each tomogram. The association of each ring subunit to individual NPCs was determined using cryoCAT package [Turoňová, B. https://github.com/turonova/cryoCAT].

### NPC ring subunit interior angle of polygon (IAOP) measurements

To obtain the interior angle of the polygon (IAOP) (see Figure 5C cartoon), i.e. the rotation around the symmetry axis from a given to the neighboring subunit, in each NPC ring (CR, IR, NR) an in-house python script was used. First, the circle center for each NPC ring was determined using preexisting code [https://meshlogic.github.io/posts/jupyter/curve-fitting/fitting-a-circle-to-cluster-of-3d-points/]. Here a threshold of at least four subunits per NPC ring was used to ensure accurate circle fitting. Then the angle between neighboring vectors connecting the circle center to the center of each subunit as determined by template matching was calculated. To ensure that only the direct neighbor angles were measured a threshold of 55° was applied and then only NPC rings with three or more angle measurements were included in the final analysis. To determine statistical significance of the measured angles in NPC rings between infected and control macrophages a threshold of 42.5° (the halfway point between the angle of a regular C8-symmetric NPC with 45° and the angle of a C9-symmetric NPC with 40°) was chosen. Then the number of NPCs that had a median subunit angle of less than 42.5° and those above or equal to the threshold were extracted from the data and subjected to a Fisher’s exact test (Prism 9 software). Data was plotted using Python (Figure 5C, S4).

### In-situ capsid modeling

A multi-step approach was employed to construct an atomic model of a complete *in situ* HIV capsid. STA for hexamers and TM for pentamers applied to a single and relatively well-resolved cone-shaped capsid yielded the centers of 141 hexamers and 8 pentamers (Figure S5A1,2). From these, the capsid surface was estimated as a convex hull (Figure S5A3) using the ArtiaX boundary method ^70^ in ChimeraX ^72^. The position of missing points in the cone lattice were then determined by oversampling random points on the surface of the convex hull and keeping points closest to the expected positions based on the translation vectors of the existing neighbors. Those with the most neighbors were iteratively selected, adding them one-by-one to the list. Once complete, the positions of the 46 added lattice points (∼24% of the total) were visually inspected and manually adjusted in ChimeraX ^72^ (Figure S5A4).

In the next step, the lattice was globally relaxed by annealing a coarse-grained particle model (Figure S5A5). Hexamers and pentamers were represented as beads. The six neighboring beads of hexamers, and the five of pentamers were connected by harmonic springs *κ*(*r* – *r*_0_)^2^/2 with *r* the pair distance, *r*_0_ = 9.3 nm the distance between hexamers obtained by placing atomic models ^74^,^75^ in the experimental STA of the hexamer which included the first neighbor, and *κ* = 3 kJ⋅mol^-1^⋅nm^-2^ the spring constant. In addition, the beads were softly tethered to their initial position by a potential *kd*^2^/2 with *d* the distance and *k* = 1.0 kJ⋅mol^-1^⋅nm^-2^ the spring constant. The total energy was annealed using Gromacs 2022 ^76^ by reducing the temperature linearly from 300 K to 10 K in 10000 steps of molecular dynamics.

Finally, the complete capsid was built by replacing each bead with atomic models of hexamers (PDB ID: 8ckv ^74^) and pentamers (PDB ID: 8g6l ^75^) rotated and translated according to the lattice (Figure S5A6). Missing loop residues were modeled using Swiss model ^77^.

### Coarse-grained model of NPC and HIV capsid

We built coarse-grained models of the NPC and the HIV capsid in which each amino acid of the proteins is mapped into a single particle ^44^. The beads are categorized as protein residues (p), membrane particles (m) and HIV capsid particles (c). The protein group has further sub-categories: scaffold residues (sc) and FG residues (FG) of NPC proteins; outer (CA) and inner (CA^i^) residues of CA hexamers and CA pentamers building the HIV capsid; and the particles containing inside the HIV capsid (HIV_in_). The potential energy of the system was given by

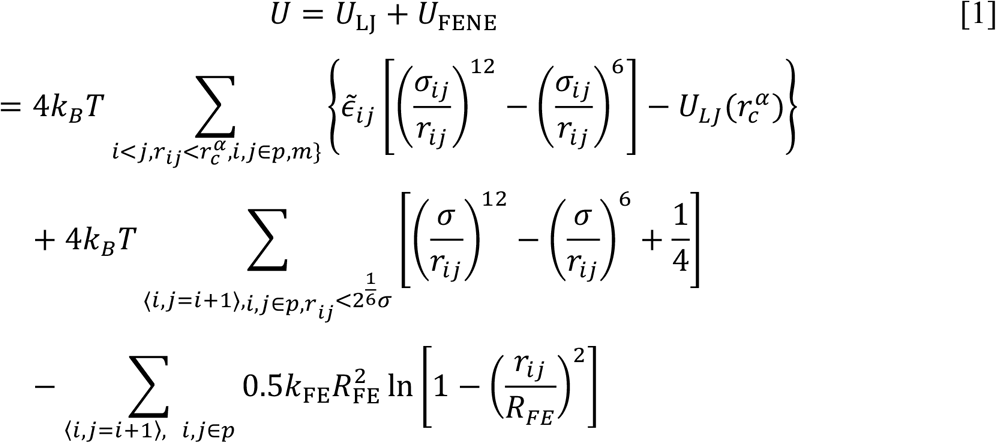

The non-bonded interactions between particles *i* and *j* in all categories are modelled by Lennard-Jones (LJ) potentials, whose strength 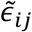, length *σ*_*ij*_, and cut-off values 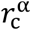 are listed in Table S2A. The bond potential between neighboring beads along the FG-proteins is expressed by the FENE potential ^78^ with *k*_*FE*_ = 30*k*_*B*_*T* and *R*_*FE*_ = 1.5*σ*. All simulations were performed in LAMMPS (Release date: Sept. 2021) ^79^. Times are reported in units of 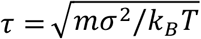, with *m* the bead mass. Box sizes are listed in Table S2B, and MD run lengths in Table S2C.

A coarse-grained representation of the nuclear envelope separating the cytoplasm and nucleoplasm was built as described ^44^. We first built a tightly packed 100×100 nm^2^ coarse-grained POPC lipid bilayer patch (command: *insane*.*py -l POPC -x 100 -y 100 -z 100 -a 0*.*3 - o bilayer*.*gro*). With this patch, we created a half-toroidal membrane pore using the BUMPy software ^80^ (command: *bumpy*.*py -s double_bilayer_cylinder -f bilayer*.*gro -z 10 -g l_cylinder:10 r_cylinder:550 r_junction:120 l_flat:2560*). We then placed particle beads at the phosphate groups of the bilayer.

For the NPC, we used the dilated NPC based on PDB-ID: 7R5J ^26^ with the following FG-Nups anchored as in model II of ^44^: NUP54, NUP58, NUP62, NUP98, POM121, NUP214, NUP153 and NUP358. Both NPC scaffold and membrane particles were fixed during the simulations. A LJ potential with large repulsive range prevented the escape of FG-Nups into the lumen of the nuclear envelope (Table S2A). To match experiments ^44^, we set the FG-FG interaction strength to 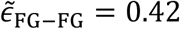 for all simulations unless otherwise stated. For 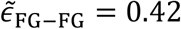, FG-Nup98 (aa1-499) was found to be close to the critical point of protein condensate formation ^44^. We confirmed that for 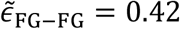 the FG-Nup98 root mean squared extension in the dilated NPC structure used here agrees with the experimental FLIM-FRET measurements (Figure S5C).

### Labelling the outer surface particles of HIV capsid

In our model, FG-Nups bind only to the outer surface of the HIV capsid. To identify the outer surface, we placed the coarse-grained HIV capsid inside a cubic box of size 441*σ* with 500 FG-Nup98 chains (aa1-499)) and ran MD simulations using a Langevin thermostat ^81^ with damping coefficient 10*τ*. We treated the HIV capsid as a rigid body using a rigid body integrator with a Langevin thermostat and a damping coefficient 3000*τ* (LAMMPS command: *fix rigid langevin molecule*). With weak interaction strengths between FG-Nup and HIV capsid particles, 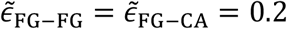, the chains formed coil configurations and explored the outer surface of HIV capsid. We did not observe the penetration of chains into the HIV capsid. Particles within 1.5*σ* of the capsid were stored every 1000*τ* for 4.5 × 10^5^*τ*. We labelled an HIV capsid particle as an outer particle if it contacted at least one FG-Nup particle during the simulation. We then labelled all equivalent particles across the 182 CA hexamers and 12 CA pentamers as outer particles, and all others as inner particles. We then filled the interior of the capsid with “cargo,” primarily RNA, modeled as particles on a cubic lattice (400807 particles with lattice constant 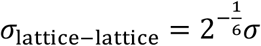). Figure S5B shows the cut view of the HIV capsid with inner lattice particles. We do not show the inner lattice particles in other figures.

### Binding affinity of a strand of FG-Nup153 to CA hexamers

We estimated the effective FG-CA hexamer interaction strength by matching the calculated binding affinity of a single CA hexamer and a fragment of FG-Nup153 to the experimentally measured value ^17^. A single CA hexamer and a 17-mer fragment of FG-Nup153 (aa1407-aa1423) were simulated inside a cubic box of size 60*σ* = 36 nm. We sampled the cross interaction potential between CA hexamer and oligomer *U*_FG−CA_(*t*) every 10*τ* for 10^7^*τ*, with pair interactions truncated and shifted at 2*σ*. We defined CA hexamer and oligomer as bound if the interacting energy was lower than the threshold energy: *U*_FG−CA_ < *U*_th_. The probability of the bound state was calculated as 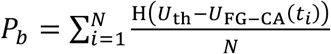, where *N* = 10^6^ is the number of sampled energies, and H(*x*) = 1 for *x* > 0 and H(*x*) = 0 otherwise is the Heaviside function. The dissociation constant as a measure of the binding affinity is calculated as 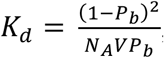, where *N*_*A*_ is the Avogadro number and *V* = (36nm)^3^ the box volume. As shown in Figure S5D,E, we found that for interaction strengths close to 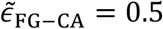 the calculated binding affinity matches the experimental value 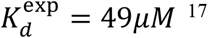 independent of threshold energy and damping coefficient of the Langevin thermostat.

### Modelling cracks in NPC scaffold

We modelled a crack in the NPC scaffold by cutting its rings and pulling them apart at the seam to resemble a nine-fold symmetric scaffold. For this, the coordinates of scaffold beads in the cylindrical coordinate system {r, *θ*, z} were mapped into new coordinates {r_n_, *θ*_n_, z_n_} according to

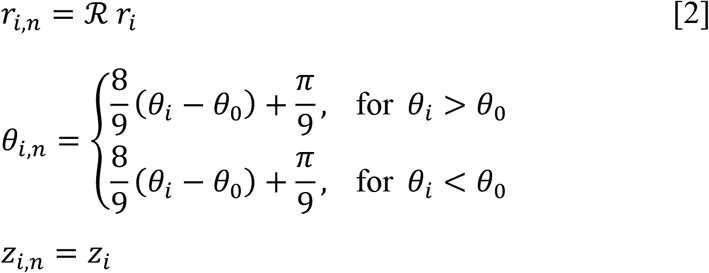

ℛ is the radial scaling factor, *θ*_0_ is the reference angle at which the NPC scaffold is cracked and the z coordinates of the beads are kept identical as in the intact NPC. This map retains the local interactions of the structure except at the crack. For scaling factors of ℛ = 1.2 and ℛ = 1.45, the inner-ring diameters *D*_in_ widens from 58 nm to 69.6 nm and 84.1 nm, respectively. With *θ*_0_ = 0.1031 rad, we set the crack interface between two spokes of the NPC scaffold in our model. To fit the nuclear envelope around the new cracked NPC, we rescaled the radial coordinates of the membrane particles by factors 1.14 and 1.38 for ℛ = 1.2 and ℛ = 1.45, respectively. The resulting structures of the NPC scaffold are shown in Figure S6A-F.

To model the dispersion in NPC diameters seen in the experiments (Figure S3D), we built intact NPC models with different scaffold pore dimensions. For this, we radially scaled the scaffold bead positions of the intact in-cell NPC without opening a lateral void by using the mapping

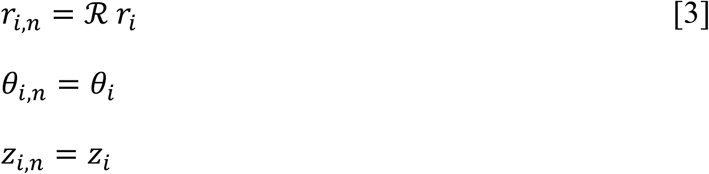

To equilibrate the cracked and intact expanded NPCs, the FENE bond potential was initially replaced by a harmonic bond potential with 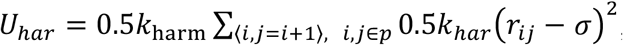, where *k*_*harm*_ = 10^4^*k*_*B*_*Tσ*^−2^ and the sum runs over all bonded neighboring beads, i.e. ⟨*i, j* = *i* + 1⟩. The systems were equilibrated for 131 × 10^3^*τ*. Then the harmonic bonds were replaced with the FENE bond and the systems were equilibrated for at least 18 × 10^3^*τ*.

### Force-driven passage of HIV capsid through NPC

We probed the passage of HIV capsids through NPCs in MD simulations with a force acting on the capsid pointing in a direction normal to the nuclear envelope. We aligned the long axis of the HIV capsid with the NPC symmetry axis. We placed the HIV capsid in the cytosol at a height of *z*_*i*_ − *z*_*c*_ = −144 nm of its center, where *z*_*i*_ is the NPC’s inner-ring position. Then we applied a force 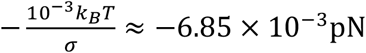 directed along the z direction toward the nucleus onto each HIV capsid particle, amounting to a total force of 4.488 nN. The total simulation runs were 10^4^ *τ*. For reference, we also simulated NPCs without FG-Nups. The position of the HIV capsid center as a function of the simulation time is shown in Figure S6G-L for three replicas. For the intact in-cell NPC (*D*_in_ = 58 nm), the HIV capsid remained stuck inside the NPC, with steric collisions blocking the translocation through the NPC scaffold even without FG-Nups. By contrast, the HIV capsid can translocate through the intact expanded NPC (*D*_in_ = 69.9 nm, ℛ = 1.2) and cracked NPCs (*D*_in_ = 69.9 nm, ℛ = 1.2 and *D*_in_ = 84.1 nm, ℛ = 1.45).

### Free energy profile for HIV capsid passage through NPC

We used MD simulations to determine the free energy profile for HIV capsid passage through NPCs. We extracted a set of initial configurations at different capsid positions from the MD simulations of force-driven NPC passage described above. We then fixed the HIV capsid in space and ran simulations for 188 × 10^3^*τ*. After 90 × 10^3^*τ* of equilibration, we averaged the total force on the HIV capsid directed along the z-axis every 10*τ*. Figures S7A-C show the final configurations for different HIV capsid positions along the translocation path. Figure S7D shows the total force on the HIV capsid per CA monomer (with a total of 1152 monomers in the HIV capsid). As described in the analysis of force-driven passage, steric clashes prevent the HIV capsid from translocating beyond *z*_*i*_ − *z*_*c*_ > −10 nm in the intact in-cell NPC. We obtained the potential of mean force by integrating the mean force along the translocation path, 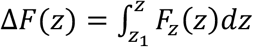, where *z*_1_ is the initial position on the cytoplasmic side. For the intact in-cell NPC, numerical integration gave a free energy minimum at *z*_*i*_ − *z*_*c*_ = −30.5 nm. For the cracked NPCs with *D*_in_ = 69.9 nm and *D*_in_ = 84.1 nm, we fitted two Gaussian functions 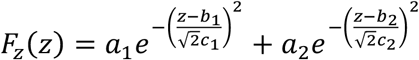 to the force data with the symmetry constraint 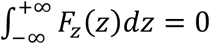. The fitting parameters are listed in Table S2D. For the cracked NPCs, the minima of the free energy are at *z*_*i*_ − *z*_*c*_ = −1 nm and 7 nm, respectively (Figure 6D).

### Release and tilt of HIV capsid inside NPC

The HIV capsid was released inside intact and cracked NPCs with 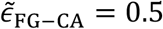. In MD simulations of at least 2 × 10^5^ (Table S2C), the capsid position and orientation was relaxed. The final orientations of the released HIV capsids are shown in Figure 6E.

## Data availability

HIV capsid hexamer and pentamer maps reported in this paper will be deposited in the EMDB with accession codes XXX and released upon publication. The NPC maps will also be deposited in the EM Data Bank with accession codes XXX and released upon publication.

The complete HIV capsid model used for simulations will be available at PDB-Dev with accession code XXX and released upon publication.

The raw tilt series and alignment files for both HIV- and mock infected conditions will be deposited on EMPIAR with accession codes XXX and available upon publication.

Initial configurations and trajectories of the MD simulations will be made available upon publication at zenodo.org under CC-BY license.

## Acknowledgements

We thank all members of the Beck, Hummer and Kräusslich laboratories for helpful discussions. We are grateful to Mark Linder and the members of the Central Electron Microscopy facility of the Max Planck Institute of Biophysics for technical support and support with data acquisition. We also thank the Infectious Diseases Imaging Platform (IDIP) at the Center for Integrative Infectious Disease Research and the Electron Microscopy Core Facilty of Heidelberg University. We are grateful to Maria Anders-Össwein, Anke-Mareil Heuser and Vera Sonntag-Buck for MDM preparation and technical support. We thank Özkan Yildiz, Juan F. Castillo Hernandez, Thomas Hoffmann, Andre Schwarz for discussions and the Max Planck Computing and Data Facility for support with scientific computing. We are grateful to Stefanie Böhm and Barbara Müller for careful reading and advice during manuscript preparation. J.P.K. thanks the International Max Planck Research School (IMPRS) on Cellular Biophysics.

## Author contributions

J.P.K., V.Z., G.H., H.-G.K. and M.B. conceived the project. V.Z. designed and performed the fluorescence microscopy and CLEM experiments with help from V.L. and analyzed data. J.P.K. performed the cryo-ET data acquisition with help from S.W.. J.P.K. performed subtomogram averaging, template matching and subsequent data analysis with help from B.T. M.H. performed the MD simulations with help from S.C.-L. The manuscript was written by J.P.K., M.H., V.Z., G.H., H.-G.K. and M.B. A.O.-K. and J.K. provided important contributions to NPC structure analysis and statistical analysis respectively. All authors contributed to manuscript editing and approved the final manuscript.

