## Supplementary Information for "Passage of the HIV capsid cracks the nuclear pore"

**A**

| # | Location at NPC | Shape | Length [nm] | Width [nm] | Comment |
|---|---|---|---|---|---|
| 1 | Cy, Docking | Cone | 106 | 61 | |
| 2 | Cy, Docking | Cone | 109 | 52 | |
| 3 | Cy, Docking | Cone | 135 | 61 | |
| 4 | Cy, Docking | Cone | 105 | 49 | Figure 3B" i |
| 5 | Cy, Docking | Cone | 98 | 55 | |
| 6 | Cy, Docking | Cone | 113 | 50 | |
| 7 | Cy, Docking | Cone | 106 | 54 | |
| 8 | Cy, Docking | Cone | 108 | 65 | |
| 9 | Cy, Docking | Cone | 107 | 62 | |
| 10 | Cy, Docking | Cone | 127 | 53 | |
| 11 | Cy, Docking | Tube | 97 | 36 | |
| 12 | Cy, Docking | Cone | 116 | 48 | |
| 13 | Cy, Docking | Cone | 97 | 58 | |
| 14 | Cy, Docking | Cone | 100 | 66 | |
| 15 | Cy, Docking | Cone | 127 | 62 | |
| 16 | Cy, Docking | Cone | 102 | 48 | |
| 17 | Cy, Docking | Cone | 107 | 53 | |
| 18 | Cy, Docking | Cone | 94 | 46 | |
| 19 | Cy, Docking | Cone | 89 | 55 | |
| 20 | Inside NPC | Cone | 105 | 55 | |
| 21 | Inside NPC | Cone | 120 | 58 | |
| 22 | Inside NPC | Cone | 92 | 44 | Figure 3A"iii |
| 23 | Inside NPC | Cone | 106 | 49 | Figure 3A" ii |
| 24 | Inside NPC | Cone | 110 | 41 | Figure 3A" i |
| 25 | Inside NPC | Cone | 117 | 49 | |
| 26 | Inside NPC | Cone | 119 | 51 | |
| 27 | Inside NPC | Cone | 119 | 57 | |
| 28 | Inside NPC | Cone | 104 | 51 | |
| 29 | Inside NPC | Cone | 128 | 57 | |
| 30 | Inside NPC | Cone | 108 | 47 | |
| 31 | Inside NPC | Cone | 132 | 59 | |
| 32 | Inside NPC | Cone | 106 | 43 | |
| 33 | Nu, basket | Cone | 103 | 51 | Figure 3C i |
| 34 | Nu, basket | Cone | 115 | 54 | |
| 35 | Nu, basket | Cone | 124 | 57 | Figure 3B" ii |
| 37 | Nu, basket | Cone | 115 | 58 | |
| 38 | Nu, basket | Cone | 114 | 61 | |
| 39 | Nu, basket | Cone | 129 | 52 | |
| 40 | Nu, basket | Cone | 110 | 46 | |
| 41 | Nu, basket | Cone | 104 | 50 | |
| 42 | Nu, basket | Tube | 93 | 49 | |
| 43 | Nu, basket | Cone | 106 | 44 | |

(Columns "Slicer view" and "Segmentation" (*xy*, *yz*) contain images.)

**B**

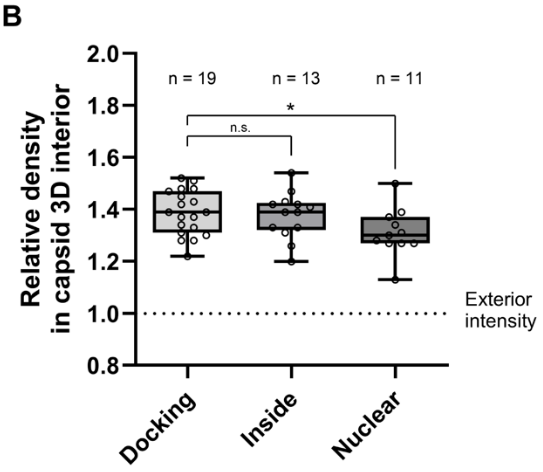


**Figure S1. Overview of viral structures captured by CLEM-ET, related to Figure 3.** (A) Capsids are shown as tomographic slices or manually segmented views (yellow). Indicated are capsids at cytoplasmic side of the NPC (Cy, docking), capsids penetrating the NPC (Inside NPC) and capsids located at basket region of the NPC (Nuclear, basket). (B) Quantification of densities observed inside of capsids captured at NPCs of infected MDM by CLEM-ET. For each of (n) segmented capsid structures the voxel intensity median within its interior was quantified and normalized to the average voxel intensity measured in the respective surrounding (dashed line) as described in Materials and Methods. Statistical significance was calculated using an unpaired two-tailed t test. n.s., not significant; *p = 0.0346. Cy, cytoplasm; Nu, nucleus.

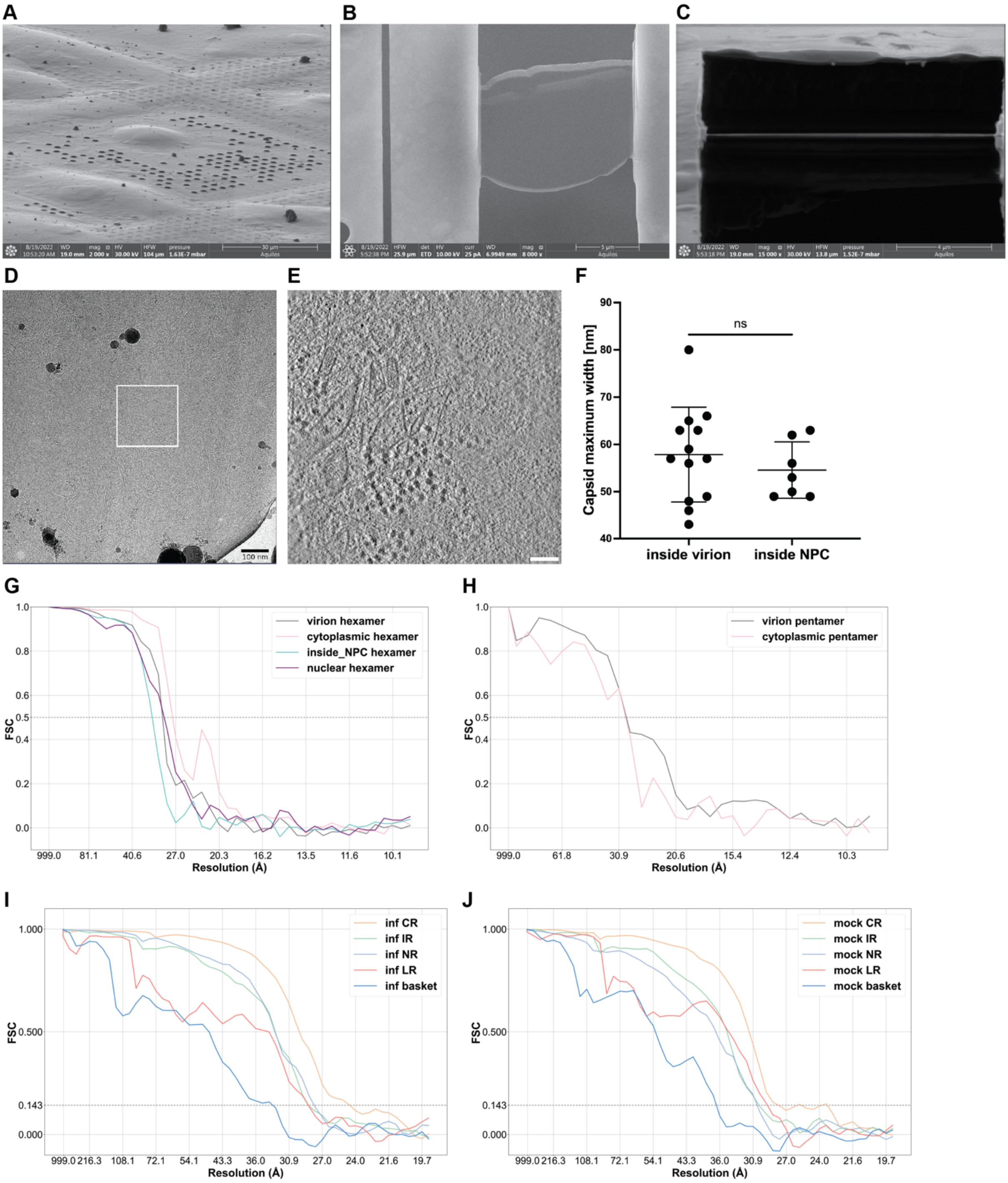


**Figure S2**. **Cryo-ET workflow of human primary macrophages, capsid widths and FSC curves, related to Figure 4.** (A) FIB view of macrophages seeded on the EM grid. (B) SEM view of thinned macrophage lamella. (C) FIB view of thinned macrophage lamella. (D) TEM medium magnification view of macrophage cell with white box highlighting acquisition area. (E) Virtual slice through exemplary cryo-tomogram. (F) Maximum capsid widths of HIV capsids inside virions and inside NPCs measured in IMOD slicer [62]. The mean width for capsids inside virions (57.9 nm, standard deviation = 10.0 nm) and the mean width for capsids inside NPCs (54.6 nm, standard deviation = 6.0 nm) are not statistically significantly different (Unpaired two-tailed t test, ns = not significant). (G) Fourier shell correlation (FSC) curves for HIV capsid hexamer STA maps. (H) Fourier shell correlation curves for HIV capsid pentamer STA maps. (I) Fourier shell correlation curves for NPC subunits from HIV-infected macrophages. (J) Fourier shell correlation curves for NPC subunits from mock-infected macrophages. For final particle numbers and resolutions of all maps see Table S1. Scale bar in (D) and (E): 100 nm.

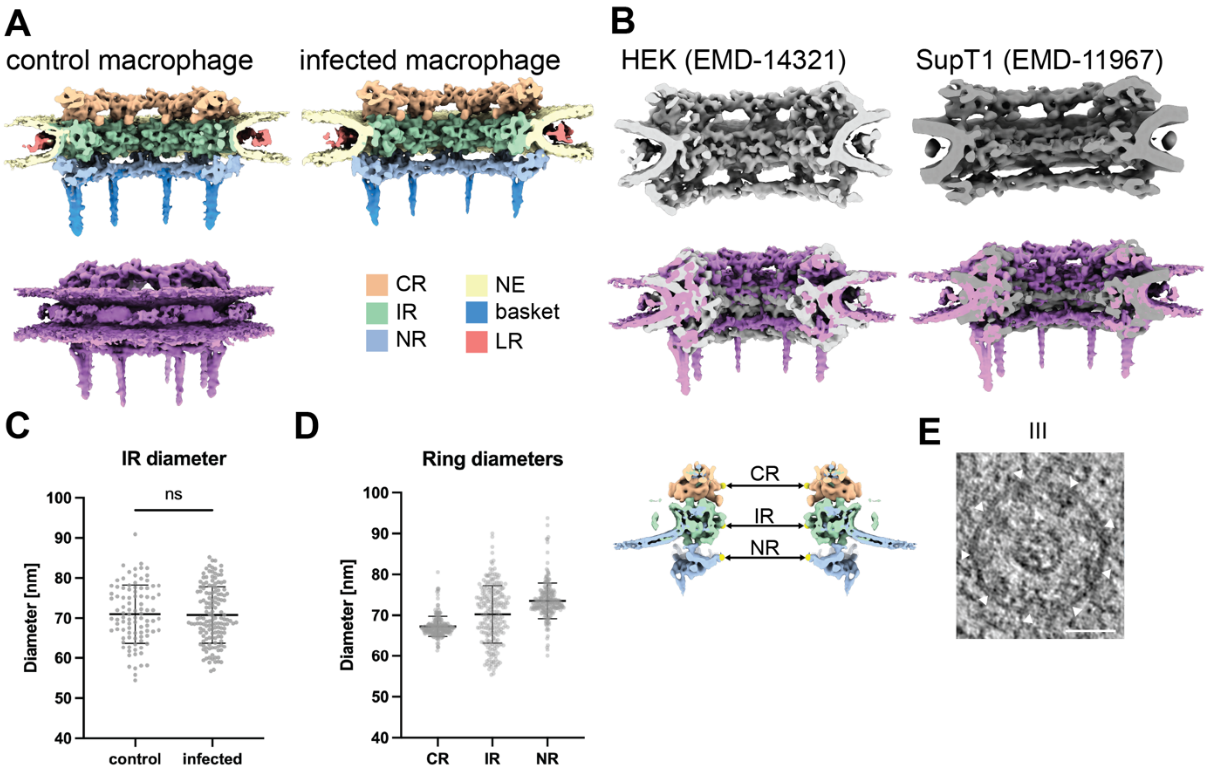


**Figure S3. Comparison of NPC structure of human primary macrophages to published human NPC structures, related to Figure 5.** (A) The macrophage NPC scaffold architecture is not altered in HIV-1 infected cells as compared to control cells. A sideview cross section of the 8-fold symmetrized NPC STA composite map shows the individual rings/filaments color-coded. The control macrophage NPC is also shown as a full sideview in pink. (B) The published NPC structure from HEK cells in light grey [26] and from SupT1 cells in dark grey [23] are shown as sideview cross sections. An overlay with the control macrophage NPC structure is shown underneath each published structure. (C) The inner ring diameters of control macrophages and infected macrophages are not significantly different. The graph below depicts the mean value and the standard deviation of the measured NPC diameters for control macrophages (mean = 71.0 nm, standard deviation = 7.2 nm, n=95) and HIV-infected macrophages (mean = 70.8 nm, standard deviation = 7.0 nm, n=145) as black bars with each data point shown as a grey dot. Unpaired two-tailed t test, ns = not significant. (D) The macrophage NPC has a wider diameter across all three rings (CR, IR, NR) as compared to SupT1 cells [23]. The schematic on top shows where the diameter was measured between opposing CR, IR, NR subunits respectively. The graphs below depict the mean value and the standard deviation of the measured NPC diameters for combined macrophages as black bars with each data point shown as a grey dot (CR mean diameter = 66.6 nm, CR standard deviation = 2.3 nm, n = 257; IR mean diameter = 69.5 nm, IR standard deviation = 7.6 nm, n = 247; NR mean diameter = 72.9 nm, NR standard deviation = 3.0 nm, n = 224). (E) For the capsid containing NPC III from Figure 5, an additional computational slice through the NPC in the tomogram is shown with white triangles highlighting the IR subunits (scale bar: 50 nm).

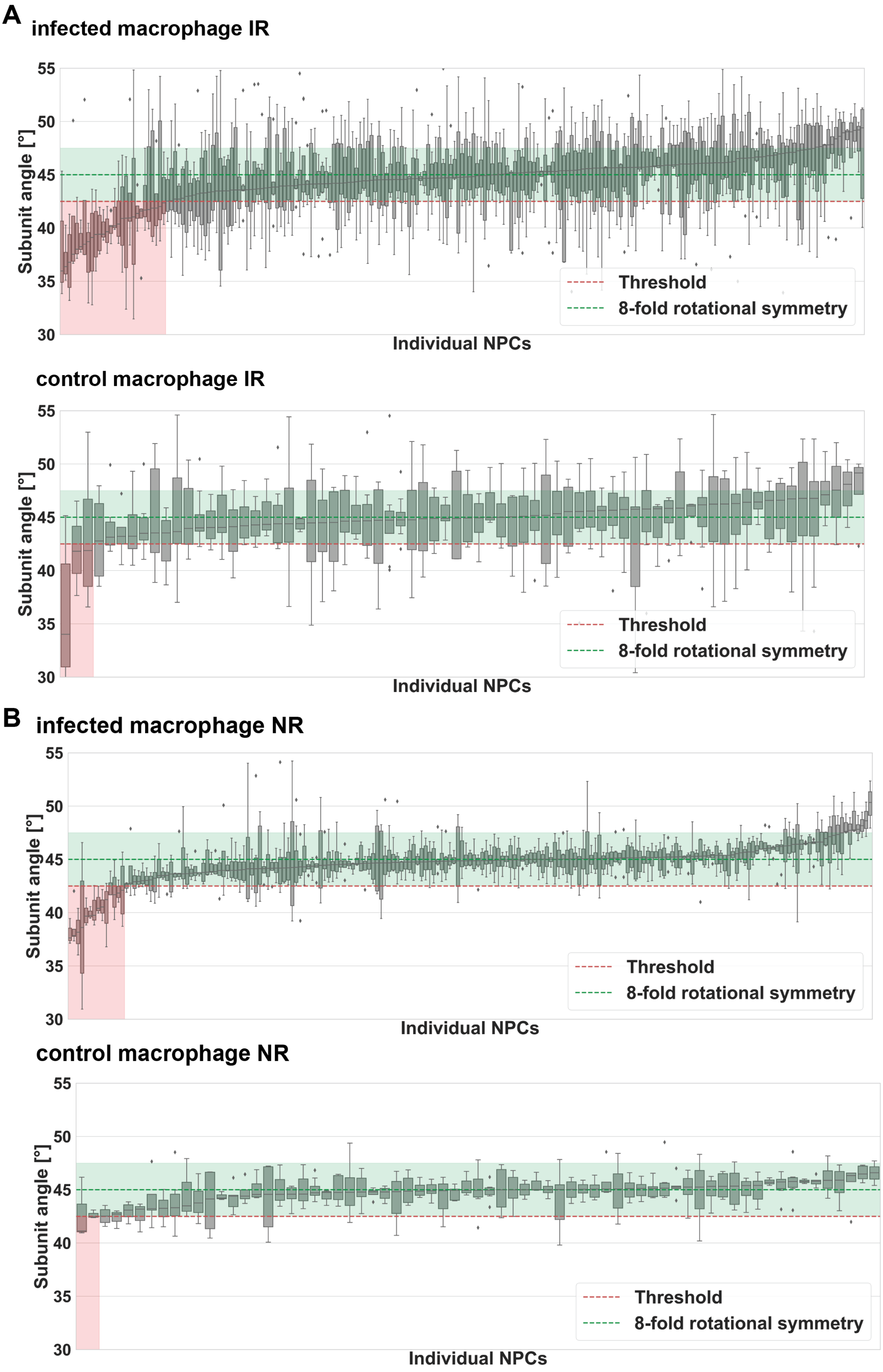
A
infected macrophage IR
Subunit angle [°]
55
50
45
40
35
30
Threshold
8-fold rotational symmetry
Individual NPCs
control macrophage IR
Subunit angle [°]
55
50
45
40
35
30
Threshold
8-fold rotational symmetry
Individual NPCs
B
infected macrophage NR
Subunit angle [°]
55
50
45
40
35
30
Threshold
8-fold rotational symmetry
Individual NPCs
control macrophage NR
Subunit angle [°]
55
50
45
40
35
30
Threshold
8-fold rotational symmetry
Individual NPCs

**Figure S4. Comparison of IAOP per NPC for the IR and NR of HIV-infected and control macrophages, related to Figure 5.** The IAOP (see methods) for each ring are plotted as boxplots and the rings are sorted by median IAOP. A threshold of 42.5° was chosen for the median IAOP und rings with a median below that value are shown in red transparent box. An expected range of subunit angles for regular C8-symmetric rings is shown in green transparent box (42.5° - 47.5°). (A) For infected macrophage IRs there were 185 C8-symmetric and 28 cracked open rings and for control macrophages 69 C8-symmetric and three cracked open rings (Fisher's exact test, $p = 0.0464$). (B) For infected macrophage NRs there were 185 C8-symmetric and 14 cracked open rings and for control macrophages 67 C8-symmetric and two cracked open rings (Fisher's exact test, $p = 0.3746$).

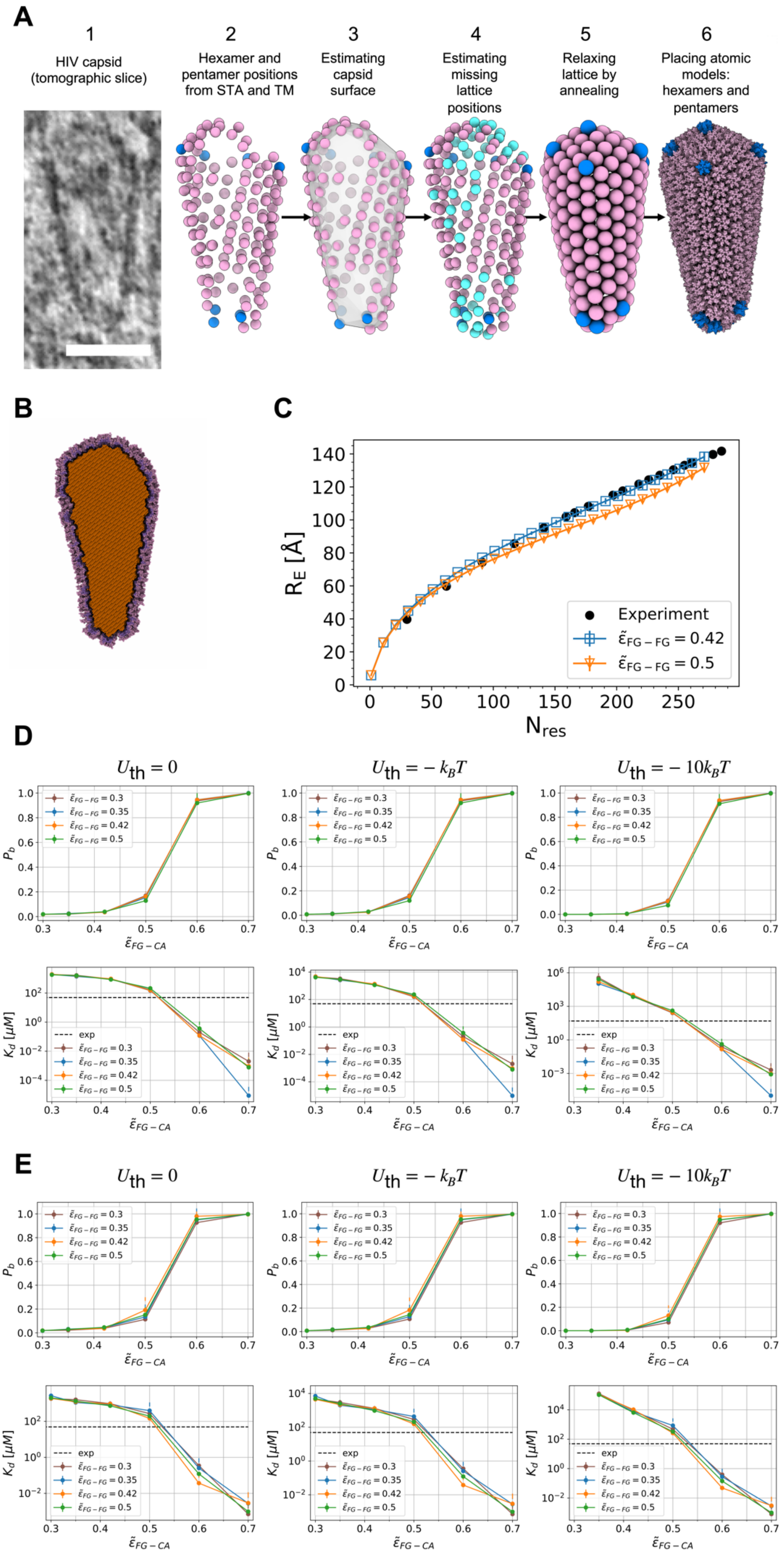

A
1
HIV capsid (tomographic slice)
2
Hexamer and pentamer positions from STA and TM
3
Estimating capsid surface
4
Estimating missing lattice positions
5
Relaxing lattice by annealing
6
Placing atomic models: hexamers and pentamers
B
C
$R_E$ [Å]
$N_{res}$
Experiment
$\tilde{\varepsilon}_{FG-FG} = 0.42$
$\tilde{\varepsilon}_{FG-FG} = 0.5$
D
$U_{th} = 0$
$U_{th} = -k_BT$
$U_{th} = -10k_BT$
$P_b$
$K_d$ [$\mu M$]
$\tilde{\varepsilon}_{FG-CA}$
exp
$\tilde{\varepsilon}_{FG-FG} = 0.3$
$\tilde{\varepsilon}_{FG-FG} = 0.35$
$\tilde{\varepsilon}_{FG-FG} = 0.42$
$\tilde{\varepsilon}_{FG-FG} = 0.5$
E
$U_{th} = 0$
$U_{th} = -k_BT$
$U_{th} = -10k_BT$

**Figure S5. Modelling of HIV-1 capsid from *in situ* cryo-ET data, related to Figure 6.**
(A) 1: Tomographic slice of exemplary cone-shaped capsid (white scale bar: 50 nm). 2: The starting positions of the majority of hexamers (pink) were obtained from STA, and the starting positions of the majority of pentamers (blue) from TM (see methods). 3: The capsid surface was estimated with the ArtiaX [70] boundary model fit. 4: The positions of the missing hexamers and pentamers (turquoise) were estimated in an iterative process (see methods). 5: The lattice was relaxed by annealing a coarse-grained particle model. 6: Atomic models of hexamers (PDB 8ckv [74]) and pentamers (PDB 8g6l [75]) were placed according to the coarse-grained lattice.
(B) Cut view of HIV capsid filled with lattice particles (orange) ordered on a cubic lattice. The interacting outer surface is shown in dark violet and the inner surface in light violet.
(C) Root-mean-squared (RMS) extension of FG-Nup98 chains inside dilated NPC as a function of separation along the sequence. For interaction strength $\tilde{\epsilon}_{\mathrm{FG-FG}} = 0.42$ (blue), the RMS extension of FG-Nup98 chains agrees with the FLIM-FRET distance measurements shown as black spheres [44]. For $\tilde{\epsilon}_{\mathrm{FG-FG}} = 0.5$ (orange), the chains are too compact. The symbols and error bars show the average and SEM estimated from four blocks of an MD simulation of $88 \times 10^3\tau$ duration.
(D, E) Binding probability $P_b$ (top) of FG-Nup153(1407-1423) to CA hexamers and the corresponding dissociation constant $K_d$ (bottom) as functions of their cross-interaction strength $\tilde{\epsilon}_{\mathrm{FG-CA}}$ for different values of the FG-FG interaction strength $\tilde{\epsilon}_{\mathrm{FG-FG}}$. Results were obtained from MD simulations with Langevin damping coefficient $10\tau$ (D) and $\tau$ (E) for binding energy thresholds of $U_{\mathrm{th}} = 0$ (left column), $U_{\mathrm{th}} = -k_BT$ (middle column), and $U_{\mathrm{th}} = -10k_BT$ (right column). For $\tilde{\epsilon}_{\mathrm{FG-CA}} \approx 0.5$, the simulation binding affinity matches with the experimental value $K_d^{\mathrm{exp}} = 49\mu M$ (dashed horizontal line [17]).

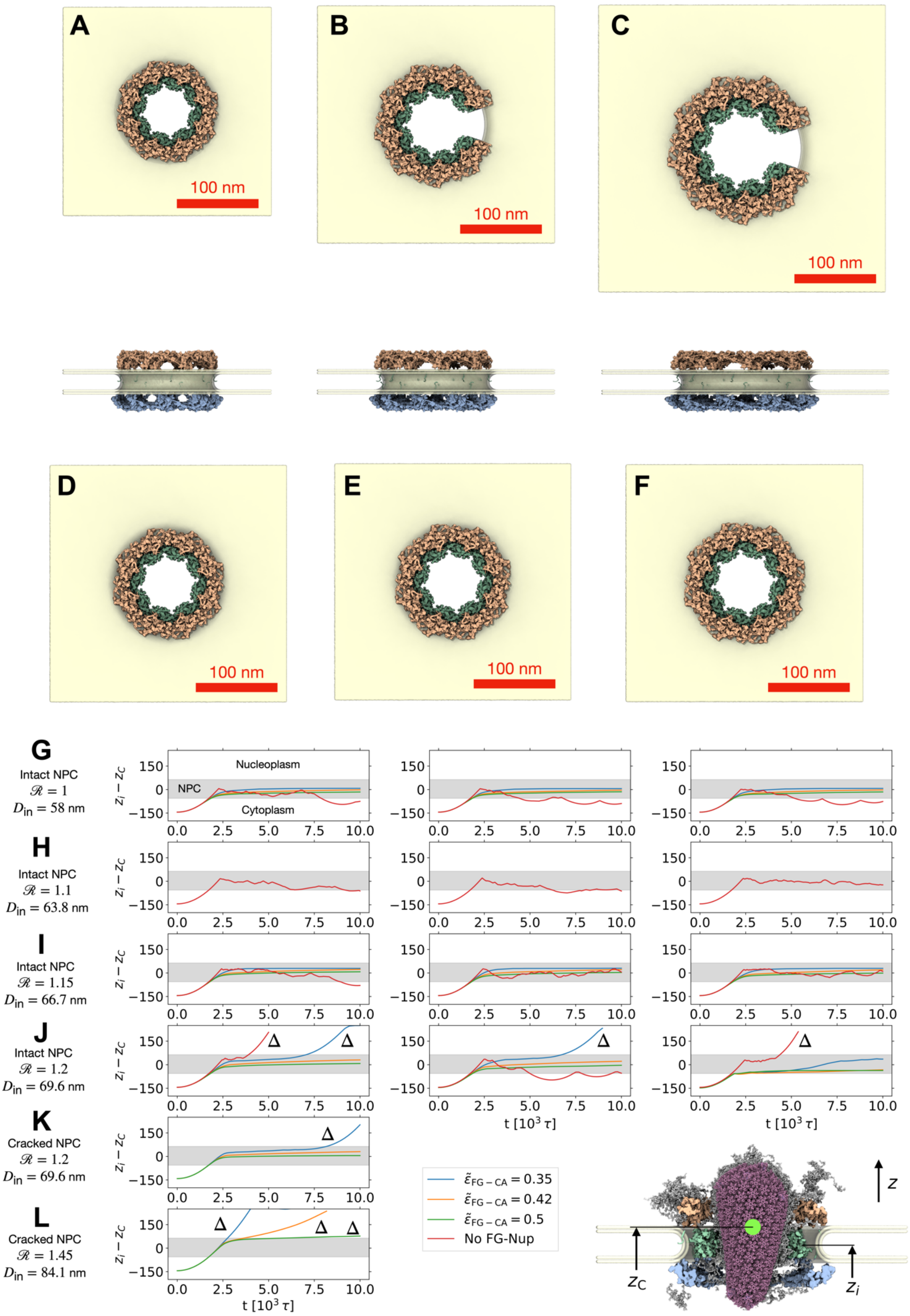
A
100 nm
B
100 nm
C
100 nm
D
100 nm
E
100 nm
F
100 nm
G
Intact NPC
$\mathcal{R} = 1$
$D_{in} = 58$ nm
Nucleoplasm
NPC
Cytoplasm
H
Intact NPC
$\mathcal{R} = 1.1$
$D_{in} = 63.8$ nm
I
Intact NPC
$\mathcal{R} = 1.15$
$D_{in} = 66.7$ nm
J
Intact NPC
$\mathcal{R} = 1.2$
$D_{in} = 69.6$ nm
K
Cracked NPC
$\mathcal{R} = 1.2$
$D_{in} = 69.6$ nm
L
Cracked NPC
$\mathcal{R} = 1.45$
$D_{in} = 84.1$ nm
$z_i - z_C$
t [$10^3\ \tau$]
$\tilde{\varepsilon}_{FG-CA} = 0.35$
$\tilde{\varepsilon}_{FG-CA} = 0.42$
$\tilde{\varepsilon}_{FG-CA} = 0.5$
No FG-Nup
$z$
$z_C$
$z_i$

**Figure S6. Passage of HIV capsid through intact, cracked and intact-dilated NPC scaffolds, related to Figure 6.** (A-F) Structure of intact NPC scaffold (A, inner ring diameter $D_{\mathrm{in}} = 58$ nm for scale factor $\mathcal{R} = 1$), cracked NPC scaffolds (B, $D_{\mathrm{in}} = 69.6$ nm, $\mathcal{R} = 1.2$; and C, $D_{\mathrm{in}} = 84.1$ nm, $\mathcal{R} = 1.45$), and intact-dilated NPC scaffolds (D, $D_{\mathrm{in}} = 63.8$ nm, $\mathcal{R} = 1.1$; E, $D_{\mathrm{in}} = 66.7$ nm, $\mathcal{R} = 1.15$; F, $D_{\mathrm{in}} = 69.6$nm, $\mathcal{R} = 1.2$) as seen from the cytosol (top) and the side (bottom). The CR, IR and NR of the NPC scaffold are shown in orange, green and blue, respectively, and the nuclear envelope in yellow. (G-L) Passage of HIV capsid through NPC scaffold in MD simulations without FG-Nups (red lines) and with FG-Nups for different interaction strengths (blue, orange, and green lines for $\tilde{\epsilon}_{\mathrm{FG-CA}} = 0.35$, 0.42, and 0.5, respectively). The HIV capsid center positions $z_i - z_C$ (inset at bottom right) are shown as function of time for the intact in-cell NPC (G, $D_{\mathrm{in}} = 58$ nm, $\mathcal{R} = 1$), the intact-dilated NPC (H, $D_{\mathrm{in}} = 63.8$ nm, $\mathcal{R} = 1.1$), the intact-dilated NPC (I, $D_{\mathrm{in}} = 66.7$ nm, $\mathcal{R} = 1.15$), the intact-dilated NPC (J, $D_{\mathrm{in}} = 69.6$ nm, $\mathcal{R} = 1.2$), the cracked NPC (K, $D_{\mathrm{in}} = 69.6$ nm, $\mathcal{R} = 1.2$) and the cracked NPC (L, $D_{\mathrm{in}} = 84.1$ nm, $\mathcal{R} = 1.45$). In (G-J), three initial conditions were tested (replicas #1-3) with the capsid rotated around its major axis by 30 degrees. In (K, L), one replica was sufficient to demonstrate that the cracked NPC scaffold is permeable to the HIV capsid. Triangles (Δ) indicate successful capsid translocation events. The position of the NPC scaffold is indicated with horizontal grey shade.

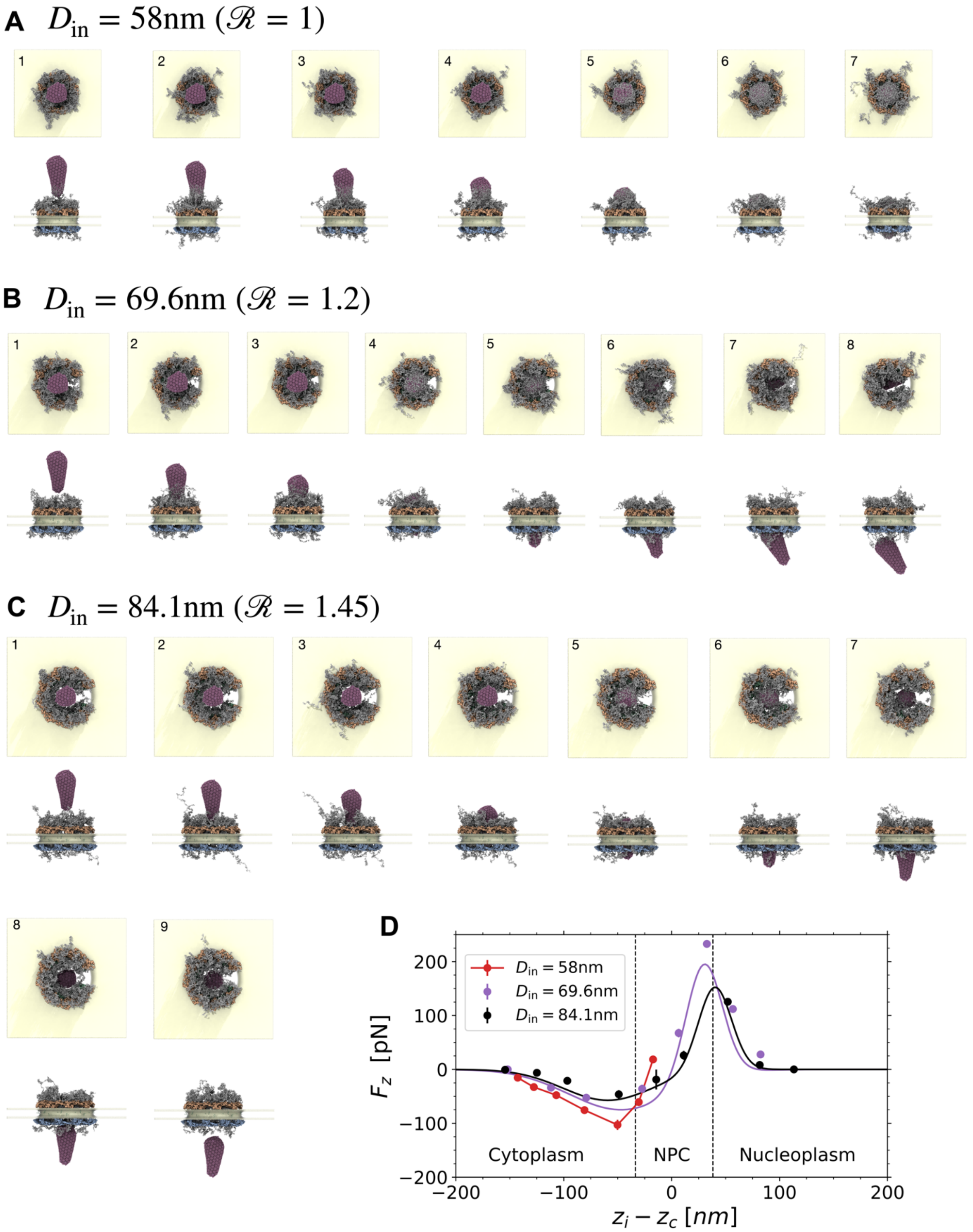


**Figure S7. Free energy of HIV capsid passage through NPCs filled with FG-Nups, related to Figure 6.**

(A) Snapshots showing final configuration in MD simulations of the intact NPC ($D_{\rm in} = 58$ nm) for different HIV capsid center positions relative to the inner ring of the NPC, $z_i - z_C = -142.7$ nm (1), $-127.7$ nm (2), $-106.9$ nm (3), $-80.8$ nm (4), $-50.2$ nm (5), $-30.5$ nm (6) and $-17.2$ nm (7). The interaction strengths were $\tilde{\epsilon}_{\rm FG-FG} = 0.42$ and $\tilde{\epsilon}_{\rm FG-CA} = 0.5$. The CR, IR and NR of the NPC scaffold are shown in orange, green and blue, respectively, the nuclear envelope in yellow, and the FG-Nups in grey.

(B) Snapshots showing final configurations of MD simulations for different HIV capsid center positions in cracked NPC ($D_{\mathrm{in}} = 69.6$ nm, $\mathcal{R} = 1.2$): $z_i - z_C = -151.9$ nm (1), $-111.8$ nm (2), $-79.1$ nm (3), $-27.3$ nm (4), 6.4 nm (5), 32.6 nm (6), 56.7 nm (7), and 82.3 nm (8). The interaction strengths were $\tilde{\epsilon}_{\mathrm{FG-FG}} = 0.42$ and $\tilde{\epsilon}_{\mathrm{FG-CA}} = 0.5$.
(C) Snapshots showing final configurations of MD simulations for different HIV capsid center positions in cracked NPC ($D_{\mathrm{in}} = 84.1$ nm, $\mathcal{R} = 1.45$): $z_i - z_C = -154.3$ nm (1), $-124.9$ nm (2), $-96.7$ nm (3), $-49$ nm (4), $-14.2$ nm (5), 11.1 nm (6), 52 nm (7), 81.5 nm (8) and 113.3 nm (9). The interaction strengths were $\tilde{\epsilon}_{\mathrm{FG-FG}} = 0.42$ and $\tilde{\epsilon}_{\mathrm{FG-CA}} = 0.5$.
(D) Mean force on HIV capsid exerted by FG-Nups and NPC scaffold. The mean force on the HIV capsid at different penetration depths $z_i - z_C$ is shown for the intact in-cell NPC scaffold ($D_{\mathrm{in}} = 58$ nm, $\mathcal{R} = 1$: red with lines as guide to the eye). For the cracked NPCs ($D_{\mathrm{in}} = 69.6$ nm, $\mathcal{R} = 1.2$: violet; $D_{\mathrm{in}} = 84.1$ nm, $\mathcal{R} = 1.45$: black), the solid lines are two-Gaussian fits with a symmetry constraint to ensure that the difference in the potential of mean force obtained by integration across the NPC is zero. For the intact in-cell NPC (red), the HIV capsid collides with the NPC scaffold, which sterically blocks passage, as indicated by the sharp rise in force. The symbols and error bars are the mean and SEM estimated from four blocks of $22.5 \times 10^3 \tau$ each. The final configurations of the systems at different positions of the HIV capsid center are shown in panels A-C. The HIV capsid was fixed during the mean-force simulations.

**Video S1**. 3D representation of an entire nuclear envelope with HIV-1 capsids, related to Figure 1D.

**Video S2**. Slices and 3D surface rendering or resin embedded tomogram, related to Figure 3A’’’.

**Video S3.** Slices through cryo electron tomogram overlaid with 3D surface rendering of ‘placed back’ NPC subunits and capsid, related to Figure 4 and 5.

**Video S4**. HIV-1 capsid lattice inside NPC, related to Figure 4 and 5.

**Video S5**. Molecular dynamics simulations of HIV-1 capsid passage through intact NPC (left) and cracked NPCs with inner-ring diameters of $D_{\mathrm{in}} = 69.6\ \mathrm{nm}$ (center) and $D_{\mathrm{in}} = 84.1\ \mathrm{nm}$ (right), related to Figure 6.

### Table S1: cryo-ET data acquisition parameters and STA map information

<table>
<tr><td>Microscope</td><td colspan="5">Titan Krios G4</td><td colspan="6">Titan Krios G4</td></tr>
<tr><td>Voltage (kV)</td><td colspan="5">300</td><td colspan="6">300</td></tr>
<tr><td>Camera</td><td colspan="5">Falcon 4</td><td colspan="6">Falcon 4</td></tr>
<tr><td>Magnification</td><td colspan="5">53000</td><td colspan="6">53000</td></tr>
<tr><td>Pixel size (Å/px)</td><td colspan="5">2.414</td><td colspan="6">2.414</td></tr>
<tr><td>Targeted total electron dose ($e^-$/Å$^2$)</td><td colspan="5">135</td><td colspan="6">135</td></tr>
<tr><td>Targeted defocus range (µm)</td><td colspan="5">-2.0 – 4.0</td><td colspan="6">-2.0 – 4.0</td></tr>
<tr><td>Automation software</td><td colspan="5">SerialEM</td><td colspan="6">SerialEM</td></tr>
<tr><td>Tomograms used for. STA/TM</td><td colspan="5">52</td><td colspan="6">97</td></tr>
<tr><td>Initial # of NPC</td><td colspan="5">121</td><td colspan="6">200</td></tr>
<tr><td>Map type</td><td>CR</td><td>IR</td><td>NR</td><td>LR</td><td>Basket</td><td>CR</td><td>IR</td><td>NR</td><td>LR</td><td colspan="2">Basket</td></tr>
<tr><td>Final # of particles</td><td>721</td><td>760</td><td>571</td><td>760</td><td>571</td><td>1185</td><td>1149</td><td>1044</td><td>1238</td><td colspan="2">1044</td></tr>
<tr><td>Resolution (Å) (FSC 0.143)</td><td>27.9</td><td>30.1</td><td>30.4</td><td>29.1</td><td>36.7</td><td>24.5</td><td>28.5</td><td>27.9</td><td>28.5</td><td colspan="2">33.2</td></tr>
<tr><td>Initial # of capsid structures</td><td colspan="5">0</td><td colspan="6">49</td></tr>
<tr><td>Map type</td><td colspan="5" rowspan="3">n.a.</td><td>Vir Hex</td><td>Cyt Hex</td><td>InNPC Hex</td><td>Nuc Hex</td><td>Vir Pent</td><td>Cyt Hex</td></tr>
<tr><td>Final # of particles</td><td>584</td><td>1675</td><td>649</td><td>585</td><td>26</td><td>58</td></tr>
<tr><td>Resolution (Å) (FSC 0.5)</td><td>30.1</td><td>27.7</td><td>33.0</td><td>29.7</td><td>29.0</td><td>29.2</td></tr>
</table>

**Table S2**

**A: Interaction parameters in coarse-grained MD simulations of HIV capsid and NPC.** The table lists the LJ interaction parameters between the different bead types. Cross-interactions parameters between beads fixed in space and beads considered as rigid body (i.e., scaffold residues, membrane beads and HIV CA hexamers and pentamers and inner particles) were not considered in the energy function.

| Type | $j \in \mathrm{sc}$ | $j \in \mathrm{FG}$ | $j \in m$ | $j \in \mathrm{CA}$ | $j \in \mathrm{CA^i}$ | $j \in \mathrm{HIV_{in}}$ |
|---|---|---|---|---|---|---|
| $i \in \mathrm{sc}$ | - | $\sigma_{ij} = \sigma$,<br>$r_c = 2\sigma$,<br>$\tilde{\epsilon}_{ij} = 0.1$ | - | $\sigma_{ij} = \sigma$,<br>$r_c = 2\sigma$,<br>$\tilde{\epsilon}_{ij} = 0.1$ | $\sigma_{ij} = \sigma$,<br>$r_c = 2\sigma$,<br>$\tilde{\epsilon}_{ij} = 0.1$ | $\sigma_{ij} = \sigma$,<br>$r_c = 2\sigma$,<br>$\tilde{\epsilon}_{ij} = 0.1$ |
| $i \in \mathrm{FG}$ | Symmetric | $\sigma_{ij} = \sigma$,<br>$r_c = 2\sigma$,<br>$\tilde{\epsilon}_{ij} = \tilde{\epsilon}_{FG-FG}$ | $\sigma_{ij} = 1.78\sigma$,<br>$r_c = 1.99\sigma$,<br>$\tilde{\epsilon}_{ij} = 0.1$ | $\sigma_{ij} = \sigma$,<br>$r_c = 2\sigma$,<br>$\tilde{\epsilon}_{ij} = \tilde{\epsilon}_{FG-CA}$ | $\sigma_{ij} = \sigma$,<br>$r_c = 2\sigma$,<br>$\tilde{\epsilon}_{ij} = 0.1$ | $\sigma_{ij} = \sigma$,<br>$r_c = 2\sigma$,<br>$\tilde{\epsilon}_{ij} = 0.1$ |
| $i \in m$ | - | Symmetric | - | $\sigma_{ij} = 1.78\sigma$,<br>$r_c = 1.99\sigma$,<br>$\tilde{\epsilon}_{ij} = 0.1$ | $\sigma_{ij} = 1.78\sigma$,<br>$r_c = 1.99\sigma$,<br>$\tilde{\epsilon}_{ij} = 0.1$ | $\sigma_{ij} = 1.78\sigma$,<br>$r_c = 1.99\sigma$,<br>$\tilde{\epsilon}_{ij} = 0.1$ |
| $i \in \mathrm{CA}$ | Symmetric | Symmetric | Symmetric | - | - | - |
| $i \in \mathrm{CA^i}$ | Symmetric | Symmetric | Symmetric | - | - | - |
| $i \in \mathrm{HIV_{in}}$ | Symmetric | Symmetric | Symmetric | - | - | - |

**B: List of simulation box sizes**.

| System | Simulation box size $(L_x \times L_y \times L_z)[\sigma^3]$ |
|---|---|
| Intact in-cell NPC model | $426.6 \times 426.6 \times 480$ |
| Cracked dilated NPC model ($\mathcal{R} = 1.2$) | $486.6 \times 486.6 \times 480$ |
| Cracked dilated NPC model ($\mathcal{R} = 1.45$) | $589.4 \times 589.4 \times 480$ |

**C: Length of simulation runs used for statistical analysis of the HIV capsid tilt angle inside NPC**.

| System | Replica #1 | Replica #2 | Replica #3 |
|---|---|---|---|
| Intact in-cell NPC model | $5 \times 10^5 \tau$ | $5 \times 10^5 \tau$ | $5 \times 10^5 \tau$ |
| Cracked dilated NPC model ($\mathcal{R} = 1.2$) | $2 \times 10^5 \tau$ | $6 \times 10^5 \tau$ | $4 \times 10^5 \tau$ |
| Cracked dilated NPC model ($\mathcal{R} = 1.45$) | $4 \times 10^5 \tau$ | $5 \times 10^5 \tau$ | $7 \times 10^5 \tau$ |

**D: Parameters obtained for two-Gaussian fits to force data**.

| System | $a_1$ | $a_2$ | $b_1$ | $b_2$ | $c_1$ | $c_2$ |
|---|---|---|---|---|---|---|
| Cracked dilated NPC model ($\mathcal{R} = 1.2$) | 216.55 | −74.86 | 29.87 | −46.84 | 16.85 | 48.8 |
| Cracked dilated NPC model ($\mathcal{R} = 1.45$) | 155.73 | −57.31 | 40.04 | −59.22 | 15.31 | 41.62 |
